# Pre-acetylation in the presence of Asf1 alters Rtt019-Vps75 selectivity: Two factor authorization: Asf1 mediates crosstalk between H3 K14 and K56 acetylation

**DOI:** 10.1101/389064

**Authors:** Joy M. Cote, Yin-Ming Kuo, Ryan A. Henry, Hataichanok Scherman, Andrew J. Andrews

## Abstract

Acetylation of histones plays a critical role in maintaining the epigenetic state of the eukaryotic cell. One such acetylation site critical for DNA damage repair is H3K56ac. In *Saccharomyces cerevisiae*, H3K56ac is thought to be driven mainly by Rtt109, a lysine acetyltransferase (KAT) that associates with the histone chaperones Vps75 and Asf1. Both of these chaperones can increase the specificity of histone acetylation by Rtt109, but neither alter the selectivity. It has been shown that histones extracted from cells (*Drosophila*), presumably containing pre-acetylated histones, can incorporate higher amounts of H3K56ac relative to recombinant non-acetylated histones. We hypothesized that histone pre-acetylation and histone chaperones could function together to drive acetylation of H3K56. In the present study, we test this hypothesis using a series of singly acetylated histones to determine the impact of crosstalk on enzyme selectivity. Our data suggest that crosstalk between acetylation sites plays a major role in altering the selectivity of Rtt109-Vps75 and that the histone chaperone Asf1 mediates this crosstalk. Specifically, we show that H3K14ac/H4 functions with Asf1 to drive H3K56ac by Rtt109-Vps75. We identified an acidic patch in Asf1 that mediates this cross-talk and show that mutations to this region can alter the Asf1 mediated crosstalk that changes Rtt109-Vps75 selectivity. These data explain the genetic link between Gcn5, which acetylates H3K14 and Rtt109. More broadly these data demonstrate that acetylation sites can dictate site selectivity even in the absence of a bromodomain and helps to explain the limited complexity that has been observed of the histone post-translational modifications patterns by global proteomic studies.

## INTRODUCTION

Lysine acetylation is a reversible post-translational modification that is known to regulate eukaryotic transcription [1, 2]. Histone acetylation functions to alter the accessibility of chromatin and provide a platform for proteins to bind, providing a crucial step in transcription, replication and DNA repair [3, 4]. One of the simplest methods by which acetylation can alter chromatin stability is to weaken the interaction between histones and DNA. One of these modifications is acetylation of lysine 56 of histone H3 (H3K56ac), which has been shown to be critical for DNA repair. This is likely due to the location of K56 in the core of histone H3 at the nucleosome dyad, with modification of this site altering DNA accessibility [5–7].

In *Saccharomyces cerevisiae*, the KAT Rtt109 (KAT11) is responsible for H3K56 acetylation, as well as H3-K9, K14, K23, and K27 acetylation [8, 9]. Rtt109 requires two structurally distinct histone chaperones [3, 10–13]. One of these histone chaperones, Vps75, is a Nap1 family member and forms a stable complex with Rtt109 [11, 14–17]. *In vivo*, Vps75 functions to stabilize Rtt109. *In vivo*, yeast Rtt109 and Vps75 are expressed in equimolar concentrations [18], implying that *in vivo* Rtt109 is stochiometrically associated with Vps75. Thus, the catalytically active complex in yeast is Rtt109-Vps75. The Rtt109-Vps75 complex is also more stable and >100-fold more active *in vitro* than Rtt109 alone [12, 13]. The second chaperone, Asf1, binds to H3/H4, splitting the tetramer complex (H3/H4)_2_ and forming a new substrate that effectively doubles the substrate concentration and increases the rate of acetylation [9]. Deletions of Rtt109 or Asf1 in yeast cells resulted in no acetylation at H3K56 and impaired DNA damage repair [18, 19]. Additionally, Vps75 and Asf1 double deletions lead to an absence of H3K9 acetylation [18]. Asf1 is capable of altering the selectivity of Rtt109 towards H3K56 *in vivo* and *in vitro*, but only if the histones used *in vitro* are extracted from eukaryotic cells and thus contain post-translational modifications (PTMs) [3, 9, 11, 13]. In the absence of PTM-histones, Asf1 increases the specificity of Rtt109-Vps75 but does not alter which residues are initially acetylated (H3K9 and H3K23) [9]. Together, these data suggest that PTMs can alter the selectivity of Rtt109-Vps75.

Recently, a crystal structure of Rtt109-Asf1-H3/H4 complex was solved for a pathogenic fungus, *Aspergillus fumigatus* (*Af*Rtt109) [20]. This structure highlights how Asf1 can alter the conformation of H3/H4. However, *Saccharomyces cerevisiae* Rtt109, the focus of this work, has been more thoroughly studied than the *A. fumigatus* system. While the overall structures have similar folds, *Af*Rtt109 is missing an insertion loop essential for Vps75 binding [14, 17], and *A. fumigatus* and *S. cerevisiae* Rtt109 have low overall sequence identity (~20%). This suggests that these two related species, although possessing homologous enzyme systems, have histone chaperone functionalities with divergent influences on specificity and selectivity. Hence, the question of how Asf1 mediates Rtt109 selectivity still remains unanswered.

In this study, we demonstrate that under conditions where we observe sequential acetylation by Rtt109-Vps75, Asf1 can drive H3K56ac after at least one round of acetylation. While no kinetic parameters can be determined from these experiments, the appearance of H3K56 acetylation later in the reaction could suggest that acetylation at some other site(s) is driving acetylation of K56. These experiments suggest that not only can H3K56ac be driven by pre-acetylation, but also that Asf1 could function with one of these acetylation sites. To date, little, if any, work has been done to assess how pre-acetylated histones impact Rtt109 selectivity, nor if histone chaperones are also influenced by PTM. The aim of this study was to determine the impact of pre-acetylated H3/H4 on Rtt109 selectivity and specificity, and moreover, to determine whether histone chaperone Asf1, in the presence of pre-acetylated H3/H4, influences sequential acetylation sites. To do this we used an orthogonal N^ε^-acetyllysyl-tRNA synthetase/tRNA_CUA_ pair in *E. coli* [21] to produce singly acetylated recombinant histones at sites that can be acetylated by Rtt109 (H3-K9, K14, K23, K27, and K56). Using these substrates, we found that every acetylation site was observed to have an impact on selectivity of Rtt109, but only H3K27ac can drive H3K56 acetylation in the absence of Asf1. In the presence of Asf1, the greatest change to selectivity came from H3K14ac, where only K56 was acetylated. Using non-acetylated H3/H4 and H3K14ac/H4 we screened mutations in an acidic patch on Asf1 to identify mutations that suppress the Asf1-H3K14ac/H4 dependent change to selectivity.

Our findings suggest that the histone chaperone Asf1 can recognize acetylated histones and alter subsequent site selectivity in the absence of a dedicated acetyl-lysine binding domain such as a bromodomain. The ability of H3K14ac to drive the selectivity to H3K56 provides a mechanistic framework to explain the genetic linkages observed between Asf1, Rtt109 and Gcn5, the acetyltransferase mainly responsible for H3K14ac *in vivo* [22–25]. Even more intriguing is the level of cross-talk between acetylation sites, which helps explain the limited complexity of the observed histone code but also begs the question: can histone acetylation function to coalesce multiple inputs from the cell to coordinate dynamic changes to chromatin? of observed rates.

## RESULTS

### Asf1 drives H3K56ac only after an initial acetylation event

Multiple labs have demonstrated the importance of Asf1 in the acetylation of H3K56 *in vivo* [3, 11, 12, 18, 19, 26], but efforts to understand this effect mechanistically have been hampered by inconsistency. Papers reporting efficient acetylation of H3K56 have either heavily relied on antibodies that have been called into question [27] or histones isolated from eukaryotic sources, which contain multiple post-translational modifications [13]. What can be agreed upon is that Asf1 increases the rate of acetylation by Rtt109-Vps75, and that Asf1 acts to split the tetramer ((H3/H4)2) into two H3/H4 dimers. In order to better understand the role of Asf1 we employed a multiplexed label-free high-throughput mass spectrometry assay [28].

Our initial experiments using recombinant histones suggested that Asf1 had no impact on driving the acetylation of H3K56 [9]. To show this, we used steady state enzyme kinetics where the concentration of Rtt109-Vps75 was kept low (E≪S) while using saturating amounts of acetyl-CoA and H3/H4 substrates, allowing us to measure the residue selectivity of acetylation. Selectivity is typically measured by the ratio of k_cat_/K_m_(s) (observed) called the specificity constant, for two different products or substrates [29]. However, we have shown that under conditions where one substrate (histone complex) is converted into multiple products (different acetylation states) and the initial rates of all products are measured simultaneously, the ratio of the observed k_cat_(s) for two residues is equal to the ratio of k_cat_/K_m_(s) observed [30]. The caveat to this approach is that it is limited to <10% of total substrate turnover and provides data only on the initial acetylation event. Therefore, if H3K56 is only acetylated after another residue, we would not observe it under these conditions. Consistent with the hypothesis that pre-acetylation may drive H3K56ac, an increase in H3K56 acetylation was only observed when we used histones isolated from chicken erythrocytes [9]. The most abundant PTM on these isolated histones is acetylation, and thus we hypothesized that acetylation may be the driving factor for H3K56ac. The most abundant sites acetylated in histones isolated from chicken erythrocytes are H3K14 with ~25% acetylation followed by K23ac at ~12%. Thus, we hypothesized that if the lysine residues are already acetylated, Rtt109-Vps75 will drive acetylation of other lysine residues found within H3.

To test this hypothesis, we used enzyme concentrations in excess of H3/H4 in the presence and absence of Asf1 (Figure 1). Normally, these conditions would only allow for one turnover but KATs are able to produce multiple products. In fact, we observe ~2.2 acetylations per histone with Rtt109-Vps75 and ~3.5 acetylations per histone in the presence of Asf1. Consistent with our hypothesis that Asf1 functions with pre-acetylation to drive H3K56ac, we observed a spike in the acetylation of H3K56 after the initial acetylation of other residues (Figure 1). Asf1 not only tripled the amount of H3K56ac, taking this PTM from the lowest abundance to the highest but it also reduces H3K9ac by more than two-fold. These data reconcile the difference between our observations and those of labs that reported enhanced acetylation of H3K56 in the presence of Asf1 [3, 11, 12, 18, 19, 26]. They do not explain how Asf1 functions with pre-acetylation to alter the apparent selectivity of Rtt109-Vps75: we hypothesize that an exosite (a secondary binding site distinct from the active site) may exist within the Rtt109-Vps75-H3/H4-Asf1 complex.

**Figure 1.**
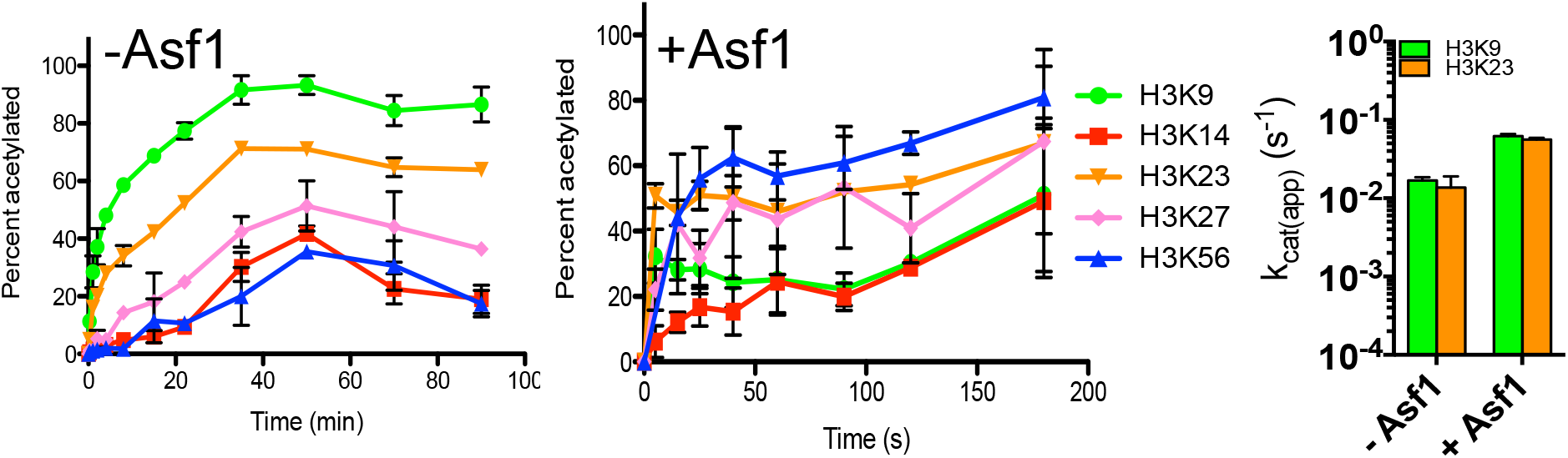
Residues acetylated within H3 by Rtt109-Vps75. A) & B) Single turnover conditions ([Rtt109-Vps75]≫[H3/H4]) with and without Asf1. C) Comparison of site-specific k_cat(app)_ of H3/H4 (with and without singly acetylated mark) by Rtt109-Vps75 acetylation in the absence or presence of Asf1. The error bar represents the standard error in k_cat(app)_. The apparent k_cat_ are summarized in Table 1 and 2.

**Table 1.** Apparent turnover numbers (k_cat_, s^-1^) of Rtt109-Vps75 for different histone substrates

| Histone Substrate | H3K9 |  | H3K14 |  | H3K23 |  | H3K27 |  | H3K56 |  |
| --- | --- | --- | --- | --- | --- | --- | --- | --- | --- | --- |
| | $k_{cat}$ (s<sup>-1</sup>) | $\Delta\Delta G^{\circ}_R$ (kcal/mol) | $k_{cat}$ (s<sup>-1</sup>) | $\Delta\Delta G^{\circ}_R$ (kcal/mol) | $k_{cat}$ (s<sup>-1</sup>) | $\Delta\Delta G^{\circ}_R$ (kcal/mol) | $k_{cat}$ (s<sup>-1</sup>) | $\Delta\Delta G^{\circ}_R$ (kcal/mol) | $k_{cat}$ (s<sup>-1</sup>) | $\Delta\Delta G^{\circ}_R$ (kcal/mol) |
| H3/H4 (WT)<sup>a</sup> | $(1.69 \pm 0.16) \times 10^{-2}$ | -3.74 | n.d. | n.d. | $(1.37 \pm 0.53) \times 10^{-2}$ | -3.64 | n.d. | n.d. | n.d. | n.d. |
| H3K9ac/H4 | n.a. | n.a. | $(6.16 \pm 0.93) \times 10^{-4}$ | -1.93 | $(1.83 \pm 0.19) \times 10^{-3}$ | -2.20 | $(4.59 \pm 1.03) \times 10^{-4}$ | -1.77 | n.d. | n.d. |
| H3K14ac/H4 | n.d. | n.d. | n.a. | n.a. | $(7.63 \pm 0.42) \times 10^{-5}$ | -0.79 | n.d. | n.d. | n.d. | n.d. |
| H3K23ac/H4 | $(1.94 \pm 0.22) \times 10^{-3}$ | -2.55 | $(5.36 \pm 0.76) \times 10^{-4}$ | -1.86 | n.a. | n.a. | $(3.23 \pm 0.21) \times 10^{-4}$ | -1.57 | n.d. | n.d. |
| H3K27ac/H4 | n.d. | n.d. | n.d. | n.d. | $(2.03 \pm 0.15) \times 10^{-4}$ | -1.32 | n.a. | n.a. | $(1.18 \pm 0.03) \times 10^{-3}$ | -2.30 |
| H3K56ac/H4 | n.d. | n.d. | $(8.42 \pm 1.16) \times 10^{-6}$ | 0.42 | $(2.82 \pm 0.38) \times 10^{-5}$ | -0.24 | n.d. | n.d. | n.a. | n.a. |

<sup>a</sup>The turnover numbers for H3/H4 (WT) were previously published and are included here for comparison.

**Table 2.**
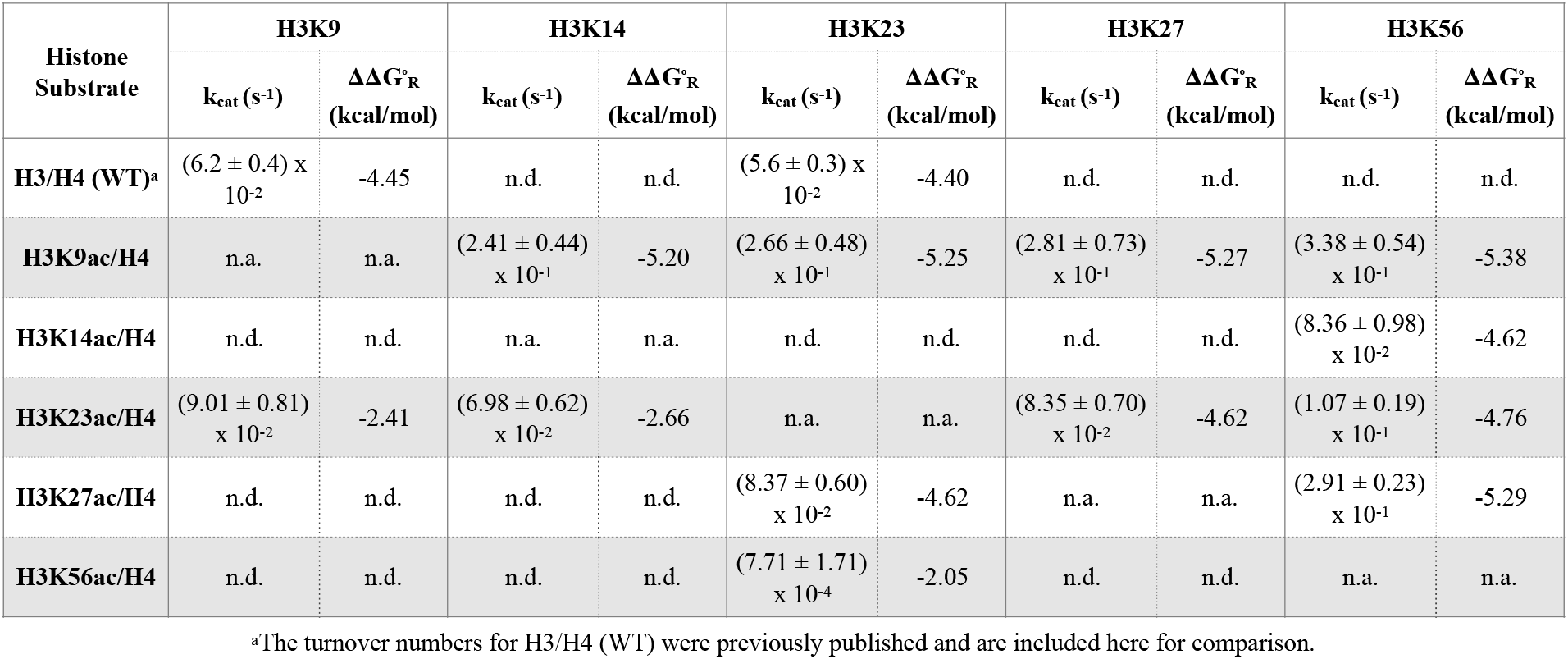
Apparent turnover numbers (k_cat_, s^-1^) of Rtt109-Vps75 for different histone substrates in the presence of Asf1

### Pre-acetylation can alter the selectivity of Rtt109-Vps75

We have now observed that pre-acetylation altered the selectivity of Rtt109-Vps75 in two experimental settings. The first, previously reported, used histones from eukaryotic cells [9] and here using an excess of enzyme over longer time courses. These experimental conditions allowed us to narrow the possible location that might alter selectivity to those residues that can be acetylated by Rtt109. To test this hypothesis, we made a library of singly acetylated histones using a strain of *E. coli* containing an orthogonal N^ε^-acetyllysyl-tRNA synthetase/tRNA_CUA_ pair [31]. The primary acetylation sites on H3/H4 by Rtt109-Vps75 were H3K9 and H3K23 [9], and H3K9 acetylation was slightly faster than H3K23 (Figure 1, Table 1). We then used these singly acetylated sites as the initial substrates to determine what impact, if any, they had on subsequent acetylation by Rtt109-Vps75. We found when H3K9ac was used as the substrate, acetylation of H3K14, H3K23, and H3K27 was observed with H3K23 being the most significant (Figure 1, Table 1). Likewise, when H3K23ac was used as the substrate, we observed the acetylation of H3K14, H3K27 and H3K9ac, with H3K9 being the most profound (Figure 1, Table 1). Pre-acetylation of H3K14 led to acetylation of H3K23 and K27 but suppressed K9, while pre-acetylation of H3K27 produced H3K23 and H3K56 (Figure 1, Table 1). The pre-acetylation of H3K56 produced the least activity and acetylated H3K14 and H3K23.

We also examined the acetylation of a histone(s) already possessing single acetyl-lysine using excess Rtt109-Vps75. Consistent with our steady state analysis, when using either H3K9ac or H3K23ac as the initial acetylated substrate, we observed that the primary acetylation site from H3K9ac was H3K23, while the primary acetylation site from H3K23ac was H3K9 (Figure 2B & D). Consistent with our steady state analysis, we found the pre-acetylated histone that lead to the most substantial increase of H3K56 acetylation was H3K27ac (Figure 2E), which is the only singly acetylated histone among these five tested substrates that can direct Rtt109-Vps75 specificity to H3K56. Lastly, we examined how either singly acetylated H3K14 or H3K56 altered the specificity of Rtt109-Vps75. We observed minimal acetylation with both H3K14ac and H3K56ac as the pre-acetylated substrates along with a significant decrease in acetylation turnover by Rtt109-Vps75. Overall, the apparent k_cat_ of H3K14ac and H3K56ac was ~2-3 orders of magnitude less than the k_cat_ from other forms of H3/H4 (Figure 1, 2C & F, and Table 1). In addition to testing the singly acetylated histones we also tested lysine (K) to glutamine (Q) mutations which are often used to mimic acetyl-lysine, By comparing pre-acetylated histone data to H3K9Q and H3K14Q mutations it is clear that these mutations are not mimic acetylation (Figure S1 and Table S1).

**Figure 2.**
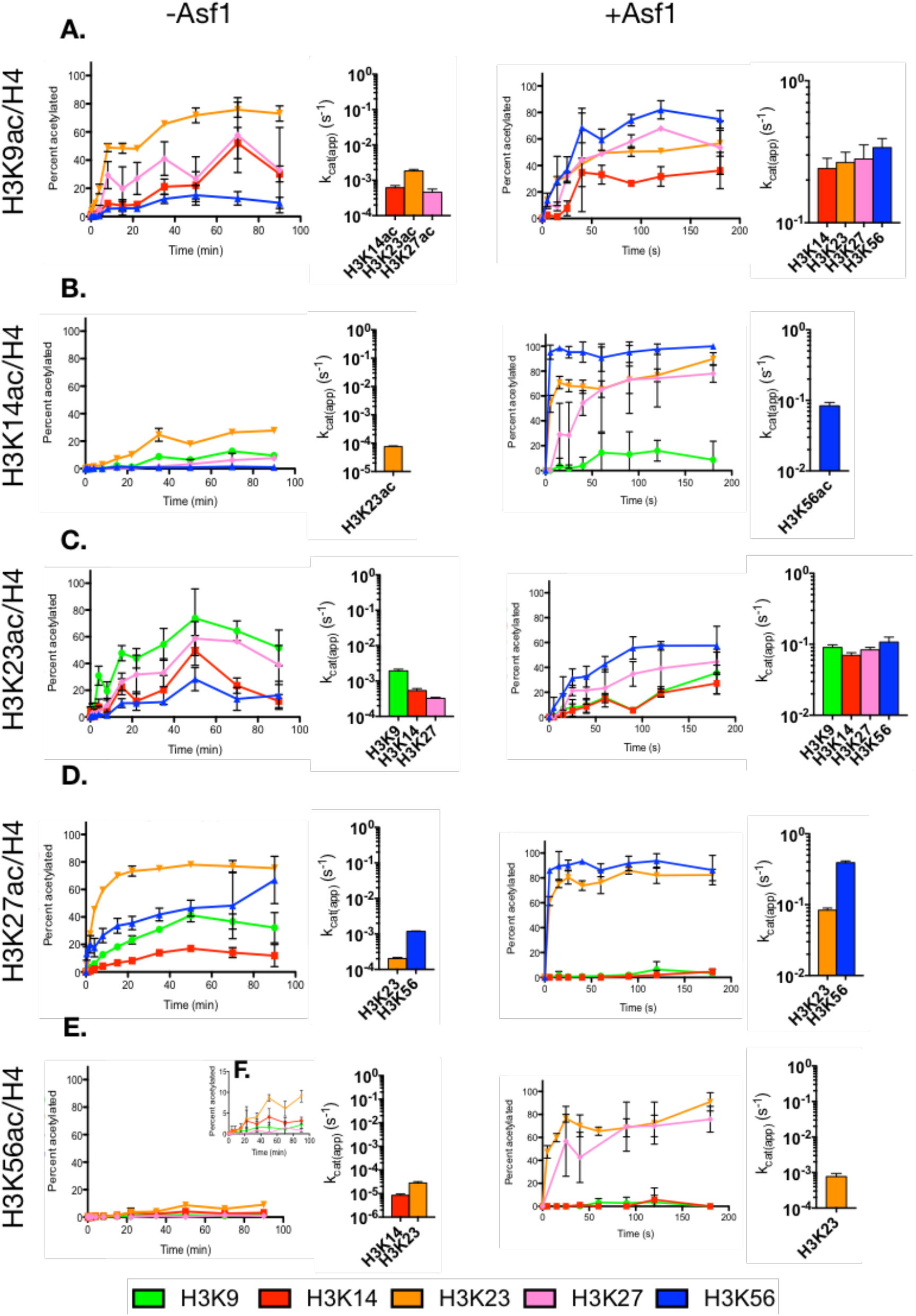
Time course site-specific plots for H3/H4 acetylation (A:WT; B: K9ac; C: K14ac; D: K23ac; E: K27ac; and F: K56ac) by Rtt109-Vps75 with and without Asf1 under single turnover and steady-state (bar graphs) conditions The individual pre-acetylated lysine's are not shown in (B-F) and remain ~100% at any time point of assays. The error bar represents the standard error in acetylation percentage. The inset in F) is to scale down the y-axis to illustrate the acetylation details of each site.

### Asf1-H3K14ac drives acetylation of H3K56ac by Rtt109-Vps75

To determine the impact of Asf1 on the residue selectivity of pre-acetylated histones we carried out experiments using limiting (steady-state) and excess Rtt109-Vps75 with various Asf1-H3KXac/H4 substrates (Figure 2). Under conditions where Rtt109-Vps75 is in excess to Asf1, several of the lysines reached a 100% acetylation in 5 seconds under the original experimental conditions (pH 7.2 and 37°C). To avoid this caveat, we used pH 6.0 and 15°C to slow down the acetylation rate, so that we would be able to differentiate the acetylation rates between sites (Figure 3). In general, the addition of Asf1 results in at least an order of magnitude increase in the apparent k_cat_ observed across all of the histone substrates and sites (Figure 3, Table 2). These results demonstrate that Asf1 can alter the selectivity of Rtt109-Vps75, but only in the presence of preexisting acetylation. Of these, H3K14ac has the largest change in selectivity, only acetylating H3K56 under steady-state conditions (~1000-fold increase). Under conditions where Rtt109-Vps75 is saturating, Asf1-H3K14ac/H4 is completely acetylated at H3K56 in <5 seconds even under low pH (6.5) and temperature (15°C) conditions.

**Figure 3.**
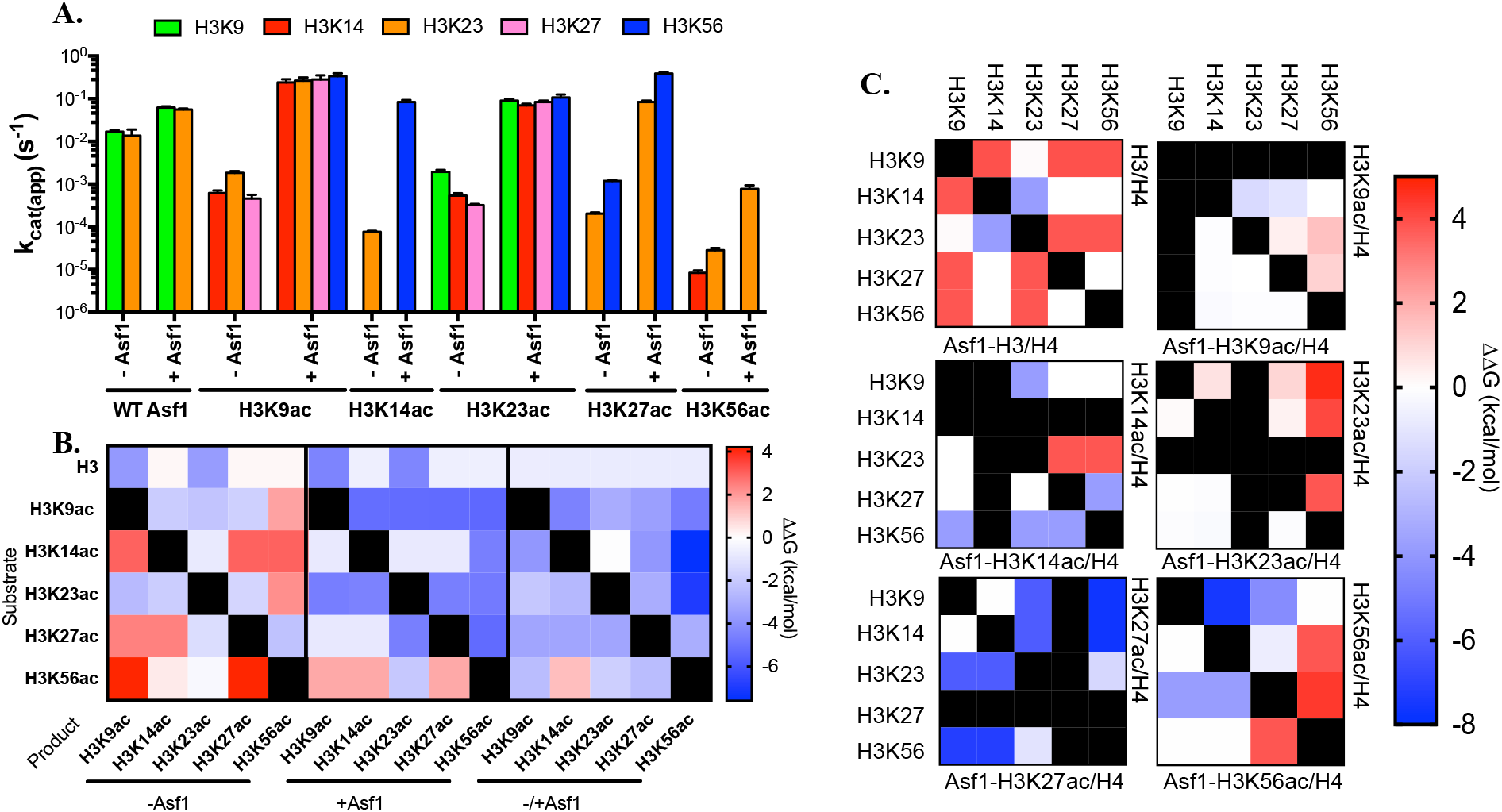
Analysis of Asf1 dependent changes to residue selectivity as a function of pre-acetylated states of histone H3. A) Comparison of site-specific k_cat(app)_ of H3/H4 (with and without singly acetylated mark) by Rtt109-Vps75 acetylation in the absence or presence of Asf1. The error bar represents the standard error in k_cat(app)_. The apparent k_cat_ are summaried in Table 1 and 2. B) Free difference for residue acetylation as compared to non-enzymatic acetylation with and without Asf1. Changes in the apparent free energy of residue selectivity due to Asf1 with varying states of histone H3. C) The apparent free energy changes to selectivity due to different acetylation states of histone H3 with and without Asf1. In these heat maps the right side of the diagonal is the selectivity between residues in the absence of Asf1 and the left side is in the presence of Asf1, and the horizontal and vertical black bars represent the sites of pre-acetylation. If Asf1 has no impact on selectivity then both sides of the diagonal will be mirror images of each other as is the case with no pre-acetylation and H3K27ac. All other changes represent changes in selectivity due to Asf1.

To confirm that we were only observing a single site of acetylation with Asf1-H3K14ac/H4, we used triton acid urea gel electrophoresis (TAU). TAU gels are able to separate different acetylation states of histones, and allow us to compare the number of acetylations per histone under different conditions. This approach confirmed that there were multiple acetylations on H3/H4 with and without Asf1, but overall acetylation increased in the presence of Asf1. H3K14ac/H4 was not turned over in the absence of Asf1, and in the presence of Asf1 we only observed a single acetylation state consistent with our MS results (Figure 4).

**Figure 4.**
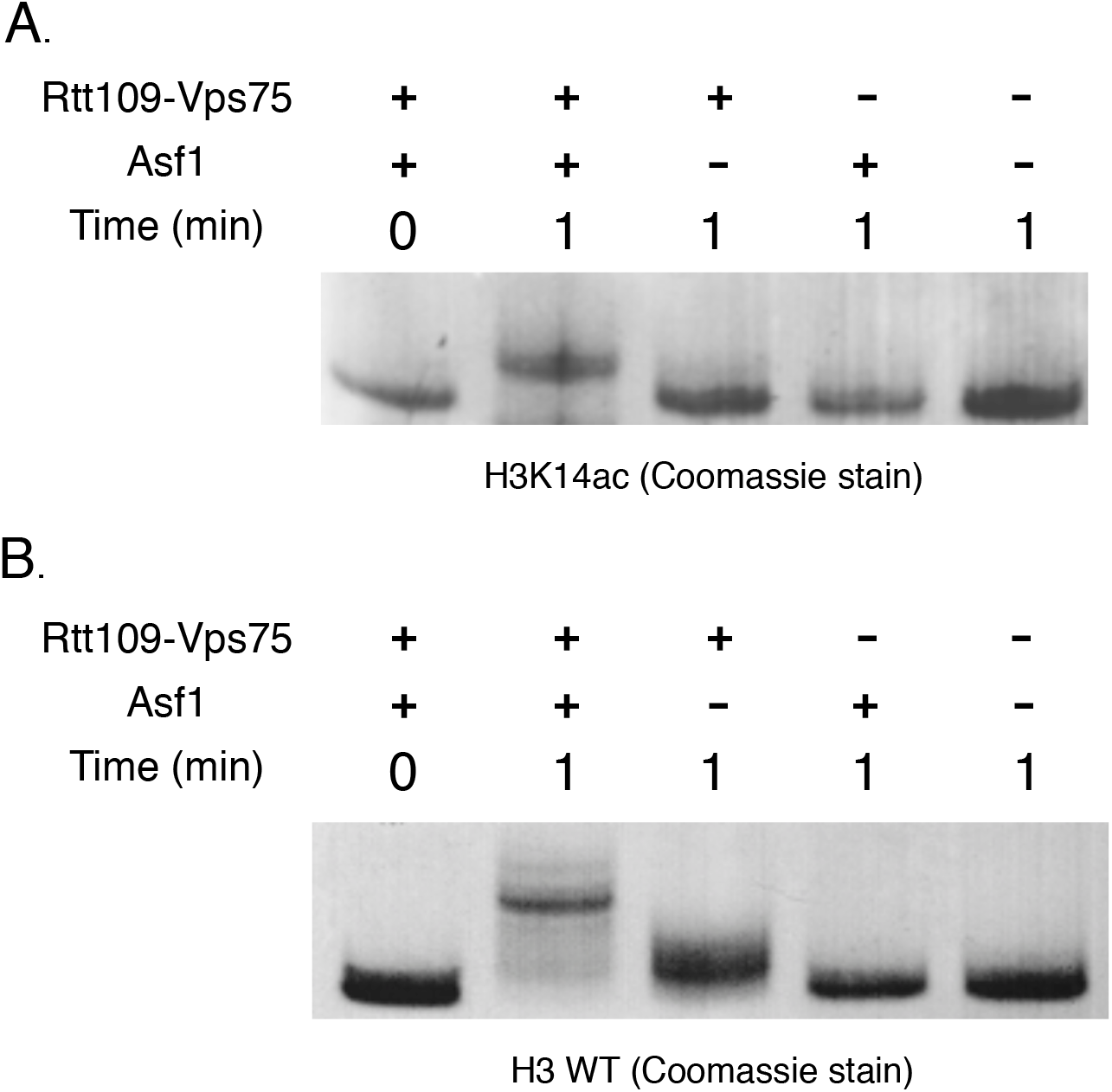
Coomassie Blue-stained triton acid-urea gel to illustrate the different acetylation states in the absence and presence of Asf1. A) H3K14ac/H4 shows low Rtt109-Vps75 acetylation activity in the absence of Asf1 but the increase of acetylation is found after Asf1 addition. B) WT H3/H4 demonstrates similar trends as H3K14ac/H4. These results are consistent with enzymatic kinetics results.

These data showed that Asf1 can increase specificity for singly acetylated histones at H3 K9, K14, K23, K27 and K56, although there was only a modest increase in specificity (27-fold increase) for H3K56ac as a substrate. For the majority of singly pre-acetylated (H3K9ac, H3K23ac and H3K27ac) substrates, Rtt109-Vps75 retained its broad specificity, acetylating multiple residues. Only H3K14ac was able to work with Asf1 to drive selectivity to exclusive H3K56 acetylation. H3K27ac was the only residue capable of driving acetylation of H3K56 in the absence of Asf1, and the only site in which we did not observe a change in selectivity, but only an increase in specificity (~200 fold).

### Comparing changes in specificity and selectivity due to pre-acetylation state and Asf1

One of the difficulties in evaluating the steady-state parameters of KATs is that many of the residues that are acetylated are acetylated at a rate too slow to be observed before 10% of the substrate has been consumed. Under these conditions we can only estimate the fastest possible rate of these residues. By comparing the fastest possible rates of these residues to the non-enzymatic rate of acetylation under the same conditions (k_nE_^200μM^) [9], we can calculate the free energy for each residue that can be acetylated. The free energy difference (ΔΔG°_R_) in the absence of Asf1 ranges from 4.2 to -3.7 kcal mol^-1^, and -0.6 to -5.4 kcal mol^-1^ with Asf1 (Figure 3B). The ΔΔG°_R_ values in the absence of Asf1 are much less favorable than in the presence of Asf1 and many are even less favorable than non-enzymatic acetylation. While Asf1 increased the favorability of acetylation, it also broadened the selectivity. There was one exception to the Asf1-dependent increase in specificity, and that was the acetylation of H3K14 (>1.3 kcal mol^-1^), which only occurred when H3K56 was pre-acetylated. The largest increase in Asf1-dependent specificity was observed with H3K14 pre-acetylation (−7.8 kcal mol^-1^), which highlights the synergy between H3K14ac and Asf1. We also compared the free energy differences in the acetylation of each site vs. every other site that can be acetylated (Figure 3C). From this analysis it becomes obvious that the Asf1 has no impact on the selectivity of H3/H4 and H3K27ac/H4, and that the pre-acetylation of K27ac increases the selectivity for both H3K23ac and H3K56ac with or without Asf1. Pre-acetylation of either H3 K9 or K23 broadened the selectivity of Rtt019-Vps75 in the presence of Asf1. Together these data demonstrate that pre-acetylation can alter the specificity and selectivity of Rtt109-Vps75, and that Asf1 can alter this pre-acetylation-dependent change in selectivity. Of all of the Asf1-dependent effects on specificity of Rtt019-Vps75 the strongest was observed with H3K14ac/H4.

### Asf1 acidic patch interacts with H3 tail to mediate acetylation

From these data, it is clear that pre-acetylation together with Asf1 facilitates a broad range of acetylation patterns. How Asf1 mediates acetylation by Rtt109-Vps75 still remains unclear. Electrostatics have been shown to be essential for many protein-protein and protein-ligand interactions. Structural analysis of the Asf1 histone complex using APBS electrostatics potential revealed an acidic patch on Asf1 (Figure 5A), that may function as an exosite that binds pre-acetylated H3 tail. Thus, we hypothesized that interactions between Asf1 and the positively charged tail of H3 is crucial for Asf1 to mediate Rtt109-Vps75 selectivity. To further investigate how Asf1 and pre-acetylated H3K14ac/H4 co-mediate H3K56 acetylation by Rtt109-Vps75, point mutations were made to individual charged residues (D37, E39, D54, D58, E88, and E105), converting each to alanine in an effort to neutralize regions of the acidic patch. Additionally, each residue was mutated to arginine to reverse the charge (Figure 5B). Steady state kinetics, as described in this report, were performed with each of the Asf1 mutants with either H3/H4 or H3K14ac/H4.

**Figure 5.**
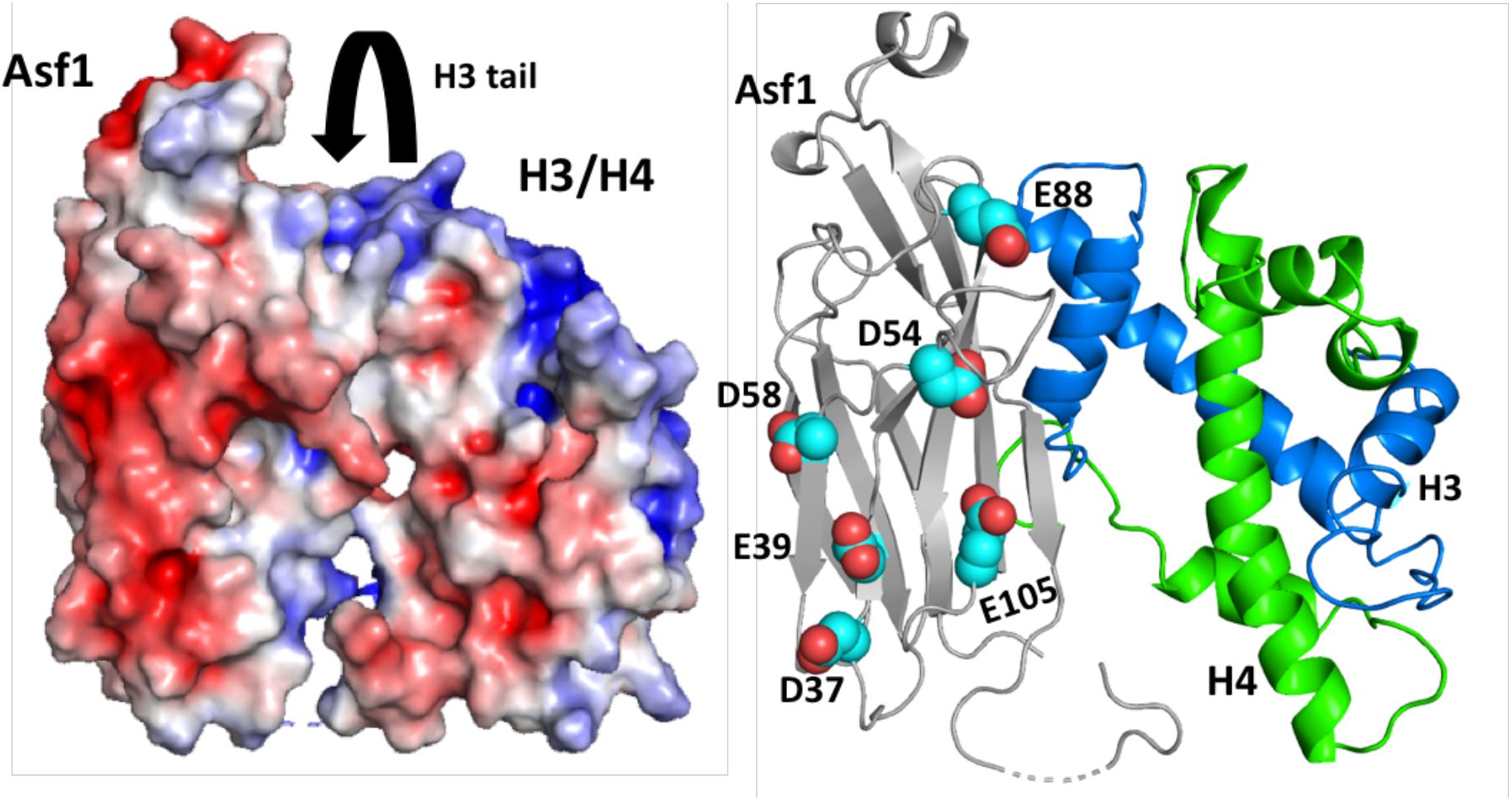
Strucutral analysis of Asf1 reveals acidic patch. A) Electronic potential surface for Asf1 H3/H4 complex without histone tail (PDB: 2io5), calculated using PyMOL plugin APBS electrostatics and default settings. B) Structure of Asf1 H3/H4 complex with proposed Asp and Glu mutation to the acidic patch highlighted in cyan.

Most Asf1 mutations altered the selectivity of Rtt109-Vps75 with H3/H4 as substrate and lead to broad acetylation at four lysine residues (Figure 6A and Table 3). However, the selectivity varied depending on the location of the mutated residue on Asf1. Alanine mutations to the top of the acidic path (D54A and E88A) lead to acetylation being observed on H3K9, H3K14, H3K23, and H3K56, with the primary acetylation site for both being H3K56. Unlike mutations to the top of Asf1, mutations to the bottom had broad selectivity. Acetylation was observed on H3K9, H3K14, H3K23 and H3K27 for mutant D37A. Similarly, for E39A acetylation was detected on H3K9, H3K14, H3K23, and H3K56. The only alanine mutant to behave like WT Asf1 was E105A. Interestingly, arginine mutants E39R, D54R, and D58R also behaved like WT Asf1 and acetylation was observed on H3K9 and H3K23. With D58A, E88R and E105R mutants, acetylation was detected at H3K9, H3K23, and H3K56, whereas D37R facilitated acetylation at H3K9, H3K14, and H3K23.

**Figure 6.**
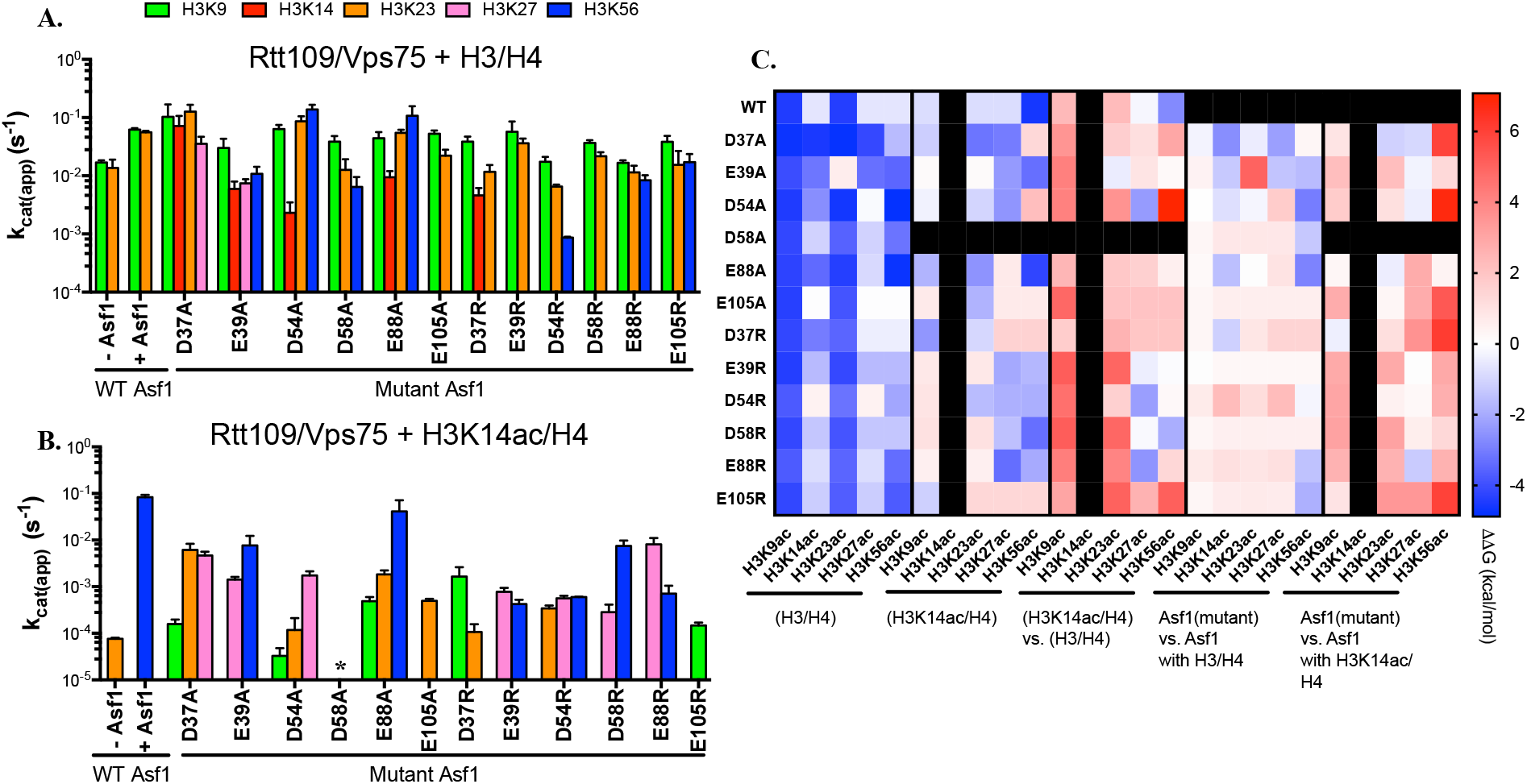
Mutants of Asf1 can suppress the Asf1 mediated cross-talk between H3K14ac and H3K56ac. Comparison of site-specific k_cat(app)_ of A) H3/H4 or B) H3K14ac/H5 by Rtt109-Vps75 acetylation in the presence of Asf1 mutants. The error bar represents the standard error in k_cat(app)_. The apparent k_cat_ are summarized in Tables 3 and 4. C) Free energy differences due to the addition of K14ac (right column) with various mutations to Asf1, and the changes to free energy due to mutations to Asf1 with H3/H4 (middle column) and H3K14ac/H4 (left column).

**Table 3.**
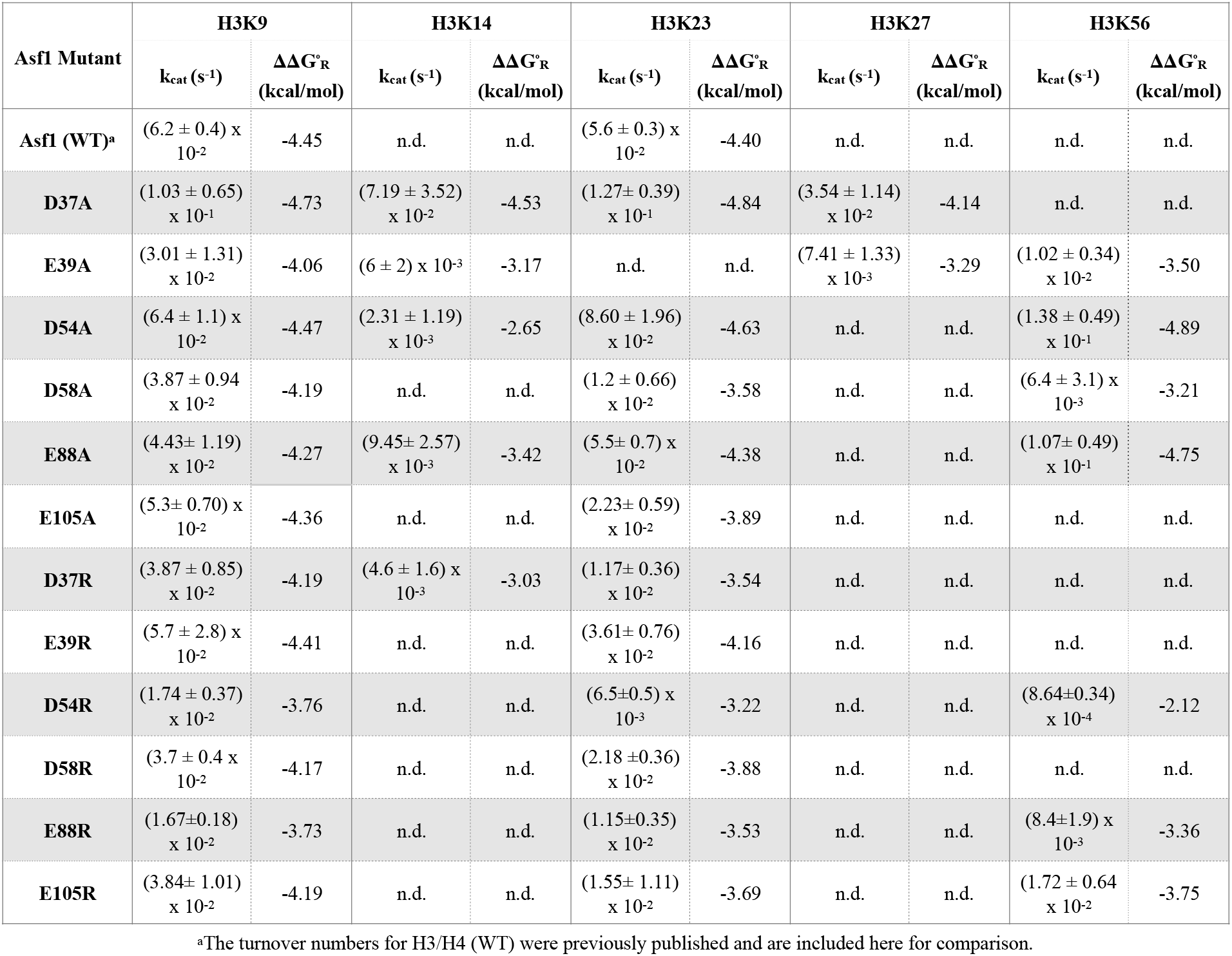
Apparent turnover numbers (k_cat_, s^-1^) of Rtt109-Vps75 for histone H3/H4 substrate in the presence of Asf1 mutants

| Asf1 Mutant | H3K9 |  | H3K14 |  | H3K23 |  | H3K27 |  | H3K56 |  |
| --- | --- | --- | --- | --- | --- | --- | --- | --- | --- | --- |
| | $k_{cat}$ ($s^{-1}$) | $\Delta\Delta G^{\circ}_R$ (kcal/mol) | $k_{cat}$ ($s^{-1}$) | $\Delta\Delta G^{\circ}_R$ (kcal/mol) | $k_{cat}$ ($s^{-1}$) | $\Delta\Delta G^{\circ}_R$ (kcal/mol) | $k_{cat}$ ($s^{-1}$) | $\Delta\Delta G^{\circ}_R$ (kcal/mol) | $k_{cat}$ ($s^{-1}$) | $\Delta\Delta G^{\circ}_R$ (kcal/mol) |
| Asf1 (WT)<sup>a</sup> | $(6.2 \pm 0.4) \times 10^{-2}$ | -4.45 | n.d. | n.d. | $(5.6 \pm 0.3) \times 10^{-2}$ | -4.40 | n.d. | n.d. | n.d. | n.d. |
| D37A | $(1.03 \pm 0.65) \times 10^{-1}$ | -4.73 | $(7.19 \pm 3.52) \times 10^{-2}$ | -4.53 | $(1.27 \pm 0.39) \times 10^{-1}$ | -4.84 | $(3.54 \pm 1.14) \times 10^{-2}$ | -4.14 | n.d. | n.d. |
| E39A | $(3.01 \pm 1.31) \times 10^{-2}$ | -4.06 | $(6 \pm 2) \times 10^{-3}$ | -3.17 | n.d. | n.d. | $(7.41 \pm 1.33) \times 10^{-3}$ | -3.29 | $(1.02 \pm 0.34) \times 10^{-2}$ | -3.50 |
| D54A | $(6.4 \pm 1.1) \times 10^{-2}$ | -4.47 | $(2.31 \pm 1.19) \times 10^{-3}$ | -2.65 | $(8.60 \pm 1.96) \times 10^{-2}$ | -4.63 | n.d. | n.d. | $(1.38 \pm 0.49) \times 10^{-1}$ | -4.89 |
| D58A | $(3.87 \pm 0.94) \times 10^{-2}$ | -4.19 | n.d. | n.d. | $(1.2 \pm 0.66) \times 10^{-2}$ | -3.58 | n.d. | n.d. | $(6.4 \pm 3.1) \times 10^{-3}$ | -3.21 |
| E88A | $(4.43 \pm 1.19) \times 10^{-2}$ | -4.27 | $(9.45 \pm 2.57) \times 10^{-3}$ | -3.42 | $(5.5 \pm 0.7) \times 10^{-2}$ | -4.38 | n.d. | n.d. | $(1.07 \pm 0.49) \times 10^{-1}$ | -4.75 |
| E105A | $(5.3 \pm 0.70) \times 10^{-2}$ | -4.36 | n.d. | n.d. | $(2.23 \pm 0.59) \times 10^{-2}$ | -3.89 | n.d. | n.d. | n.d. | n.d. |
| D37R | $(3.87 \pm 0.85) \times 10^{-2}$ | -4.19 | $(4.6 \pm 1.6) \times 10^{-3}$ | -3.03 | $(1.17 \pm 0.36) \times 10^{-2}$ | -3.54 | n.d. | n.d. | n.d. | n.d. |
| E39R | $(5.7 \pm 2.8) \times 10^{-2}$ | -4.41 | n.d. | n.d. | $(3.61 \pm 0.76) \times 10^{-2}$ | -4.16 | n.d. | n.d. | n.d. | n.d. |
| D54R | $(1.74 \pm 0.37) \times 10^{-2}$ | -3.76 | n.d. | n.d. | $(6.5 \pm 0.5) \times 10^{-3}$ | -3.22 | n.d. | n.d. | $(8.64 \pm 0.34) \times 10^{-4}$ | -2.12 |
| D58R | $(3.7 \pm 0.4) \times 10^{-2}$ | -4.17 | n.d. | n.d. | $(2.18 \pm 0.36) \times 10^{-2}$ | -3.88 | n.d. | n.d. | n.d. | n.d. |
| E88R | $(1.67 \pm 0.18) \times 10^{-2}$ | -3.73 | n.d. | n.d. | $(1.15 \pm 0.35) \times 10^{-2}$ | -3.53 | n.d. | n.d. | $(8.4 \pm 1.9) \times 10^{-3}$ | -3.36 |
| E105R | $(3.84 \pm 1.01) \times 10^{-2}$ | -4.19 | n.d. | n.d. | $(1.55 \pm 1.11) \times 10^{-2}$ | -3.69 | n.d. | n.d. | $(1.72 \pm 0.64) \times 10^{-2}$ | -3.75 |

<sup>a</sup>The turnover numbers for H3/H4 (WT) were previously published and are included here for comparison.

Each mutation utilizing H3K14ac/H4 as substrate no longer mediated acetylation solely on H3K56, as shown for WT Asf1 (Figure 6B and Table 4). Mutations to either D37A or D54A resulted in acetylation primarily being observed at H3K9, H3K23, and H327. Introducing a positive charge, however, did not lead to conserved acetylation patterns between the two mutants. D37R acetylation was observed at H3K9 and H3K23. Acetylations at H3K23, H3K27, and H3K56 were detected for D54R. Acetylations at H3K27 and H3K56 were observed for Asf1 mutants E39A, E39R, D58R, and E88R. Rtt109-Vps75 primarily acetylates H3K9, H3K23, and H3K56 with E88A mutant. Consistent with wild-type histones without Asf1, with mutant E105A only H3K23 acetylation was observed. However, upon introducing an arginine residue at E105, acetylation was observed at H3K27 and H3K56.

**Table 4.**
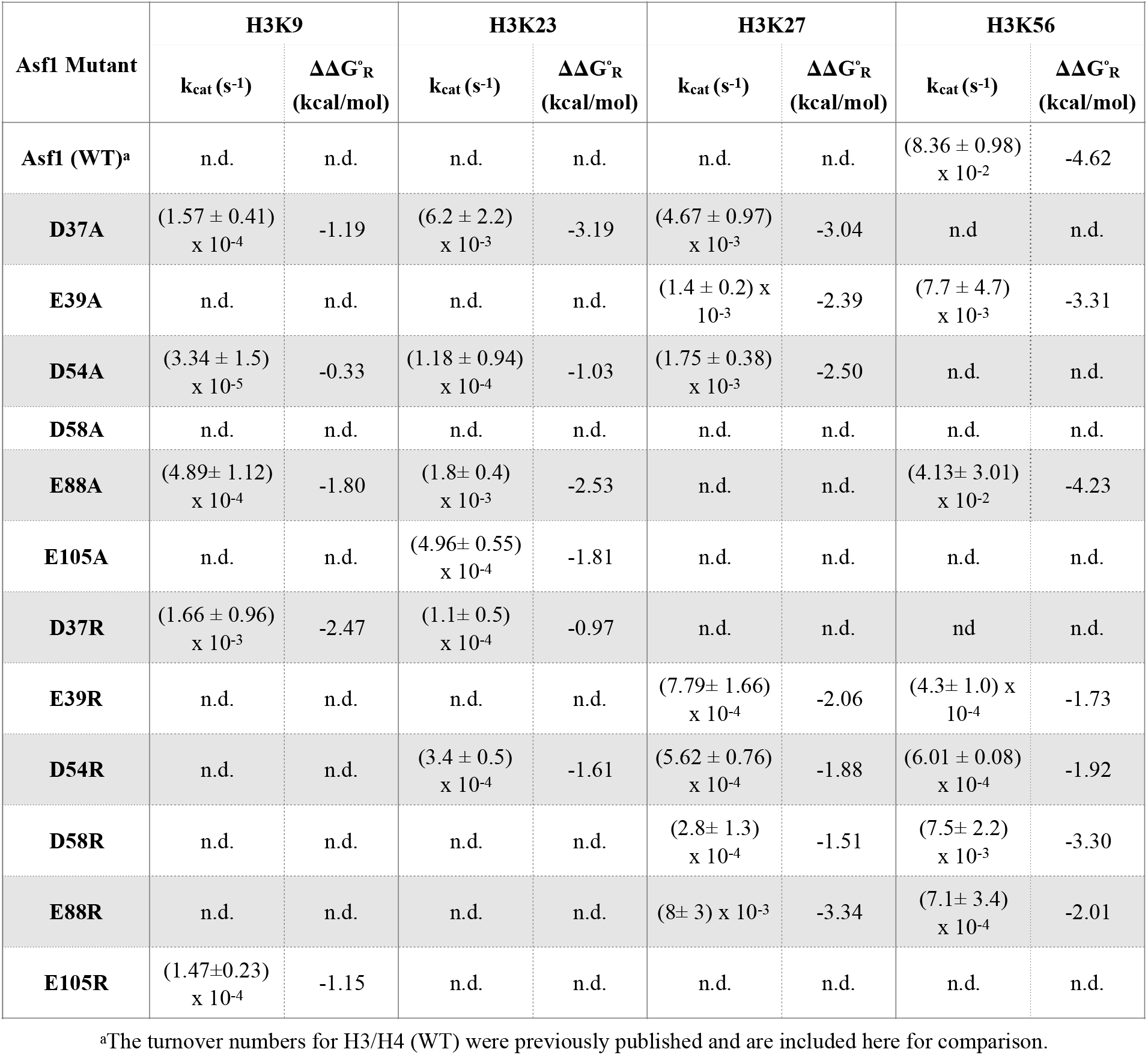
Apparent turnover numbers (k_cat_, s^-1^) of Rtt109-Vps75 for histone H3K14ac/H4 substrate in the presence of Asf1

Although all Asf1 mutations displayed interesting changes to Rtt109-Vps75 selectivity, to the E105 mutant was particularly interesting due to the loss of H3K14ac/H4 and Asf1-mediated acetylation of H3K56 with little to no changes to the acetylation of H3/H4. To further examine the impact of mutations at E105, steady state kinetics were performed with Asf1 mutants E105A/R with singly acetylated histones at H3 K9, K23, K27 and K56 (Figure 7 and Table 5). When H3K9ac/H4 was used as substrate, acetylation was observed at H3K23 and H3K56 for E105A. Interestingly, for E105R only H3K23 acetylation was detected. With pre-acetylated H3K23 and E105A, acetylation was only observed at H3K9. Conversely, E105R displayed more broad selectivity with acetylation at H3K9, H3K14, and H3K56. Like H3/H4 when H3K27ac/H4 was used as substrate with E105A, acetylation was observed at H3K9 and H3K23. Positive mutant E105R with H3K27ac/H4 displayed acetylation only at H3K14. H3K56ac/H4 was the only pre-acetylated H3 to not drive any additional acetylation with both E105A/R.

**Figure 7.**
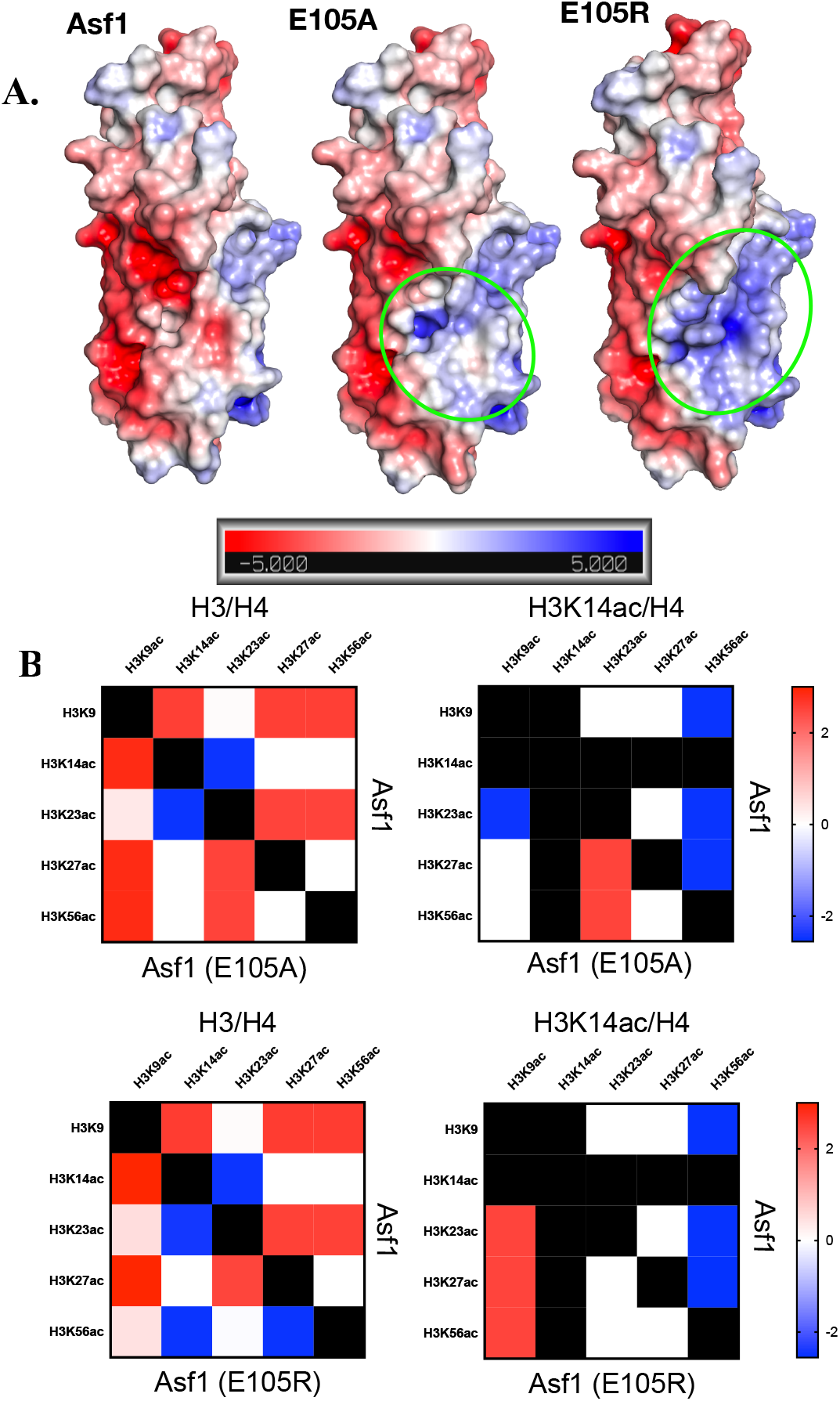
Analysis of E105 mutations impact of Asf1 acidic patch. A) Electronic potential surface for Asf1 (PDB: 2io5), calculated using PyMOL plugin APBS electrostatics and default settings. Asf1 E105A and E105R mutations were computationally made in PyMol using mutagenesis wizard and APBS electrostatics were calculated as for wild-type. B) Free energy changes to selectivity due to mutation to Asf1, where the right side of the diagonal is the selectivity between residues with wildtype Asf1 and the left side is in the presence of Asf1 with E105A or R mutations.

**Table 5.**
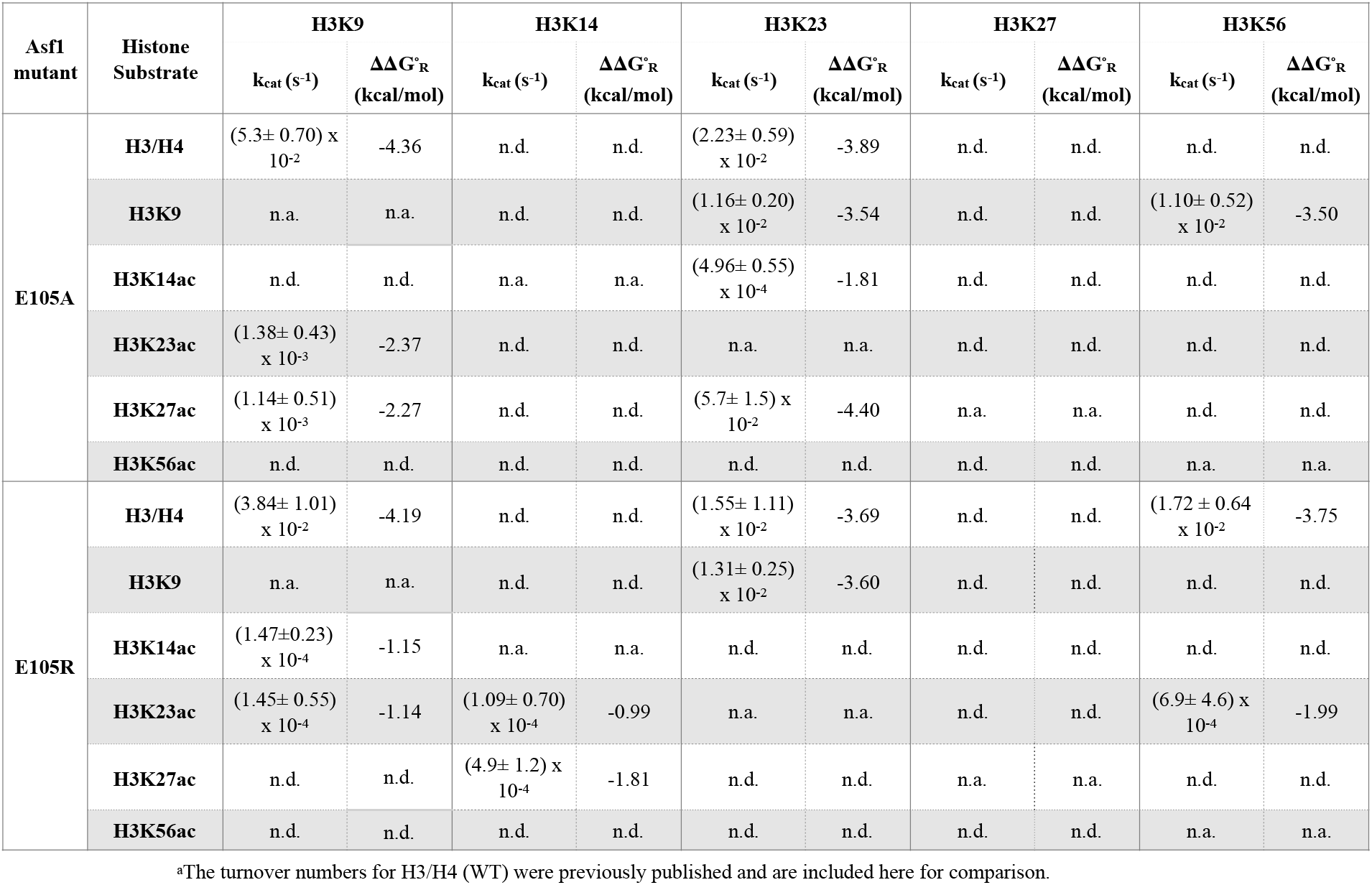
Apparent turnover numbers (k_cat_, s^-1^) of Rtt109-Vps75 for different histone substrates in the presence of Asf1 mutant E105A/R.

| Asf1 mutant | Histone Substrate | H3K9 |  | H3K14 |  | H3K23 |  | H3K27 |  | H3K56 |  |
| --- | --- | --- | --- | --- | --- | --- | --- | --- | --- | --- | --- |
| | | $k_{cat}$ (s<sup>-1</sup>) | $\Delta\Delta G^{\ddagger}_R$ (kcal/mol) | $k_{cat}$ (s<sup>-1</sup>) | $\Delta\Delta G^{\ddagger}_R$ (kcal/mol) | $k_{cat}$ (s<sup>-1</sup>) | $\Delta\Delta G^{\ddagger}_R$ (kcal/mol) | $k_{cat}$ (s<sup>-1</sup>) | $\Delta\Delta G^{\ddagger}_R$ (kcal/mol) | $k_{cat}$ (s<sup>-1</sup>) | $\Delta\Delta G^{\ddagger}_R$ (kcal/mol) |
| E105A | H3/H4 | (5.3± 0.70) x 10<sup>-2</sup> | -4.36 | n.d. | n.d. | (2.23± 0.59) x 10<sup>-2</sup> | -3.89 | n.d. | n.d. | n.d. | n.d. |
|  | H3K9 | n.a. | n.a. | n.d. | n.d. | (1.16± 0.20) x 10<sup>-2</sup> | -3.54 | n.d. | n.d. | (1.10± 0.52) x 10<sup>-2</sup> | -3.50 |
|  | H3K14ac | n.d. | n.d. | n.a. | n.a. | (4.96± 0.55) x 10<sup>-4</sup> | -1.81 | n.d. | n.d. | n.d. | n.d. |
|  | H3K23ac | (1.38± 0.43) x 10<sup>-3</sup> | -2.37 | n.d. | n.d. | n.a. | n.a. | n.d. | n.d. | n.d. | n.d. |
|  | H3K27ac | (1.14± 0.51) x 10<sup>-3</sup> | -2.27 | n.d. | n.d. | (5.7± 1.5) x 10<sup>-2</sup> | -4.40 | n.a. | n.a. | n.d. | n.d. |
|  | H3K56ac | n.d. | n.d. | n.d. | n.d. | n.d. | n.d. | n.d. | n.d. | n.a. | n.a. |
| E105R | H3/H4 | (3.84± 1.01) x 10<sup>-2</sup> | -4.19 | n.d. | n.d. | (1.55± 1.11) x 10<sup>-2</sup> | -3.69 | n.d. | n.d. | (1.72 ± 0.64) x 10<sup>-2</sup> | -3.75 |
|  | H3K9 | n.a. | n.a. | n.d. | n.d. | (1.31± 0.25) x 10<sup>-2</sup> | -3.60 | n.d. | n.d. | n.d. | n.d. |
|  | H3K14ac | (1.47±0.23) x 10<sup>-4</sup> | -1.15 | n.a. | n.a. | n.d. | n.d. | n.d. | n.d. | n.d. | n.d. |
|  | H3K23ac | (1.45± 0.55) x 10<sup>-4</sup> | -1.14 | (1.09± 0.70) x 10<sup>-4</sup> | -0.99 | n.a. | n.a. | n.d. | n.d. | (6.9± 4.6) x 10<sup>-4</sup> | -1.99 |
|  | H3K27ac | n.d. | n.d. | (4.9± 1.2) x 10<sup>-4</sup> | -1.81 | n.d. | n.d. | n.a. | n.a. | n.d. | n.d. |
|  | H3K56ac | n.d. | n.d. | n.d. | n.d. | n.d. | n.d. | n.d. | n.d. | n.a. | n.a. |

<sup>a</sup>The turnover numbers for H3/H4 (WT) were previously published and are included here for comparison.

**Figure 8.**
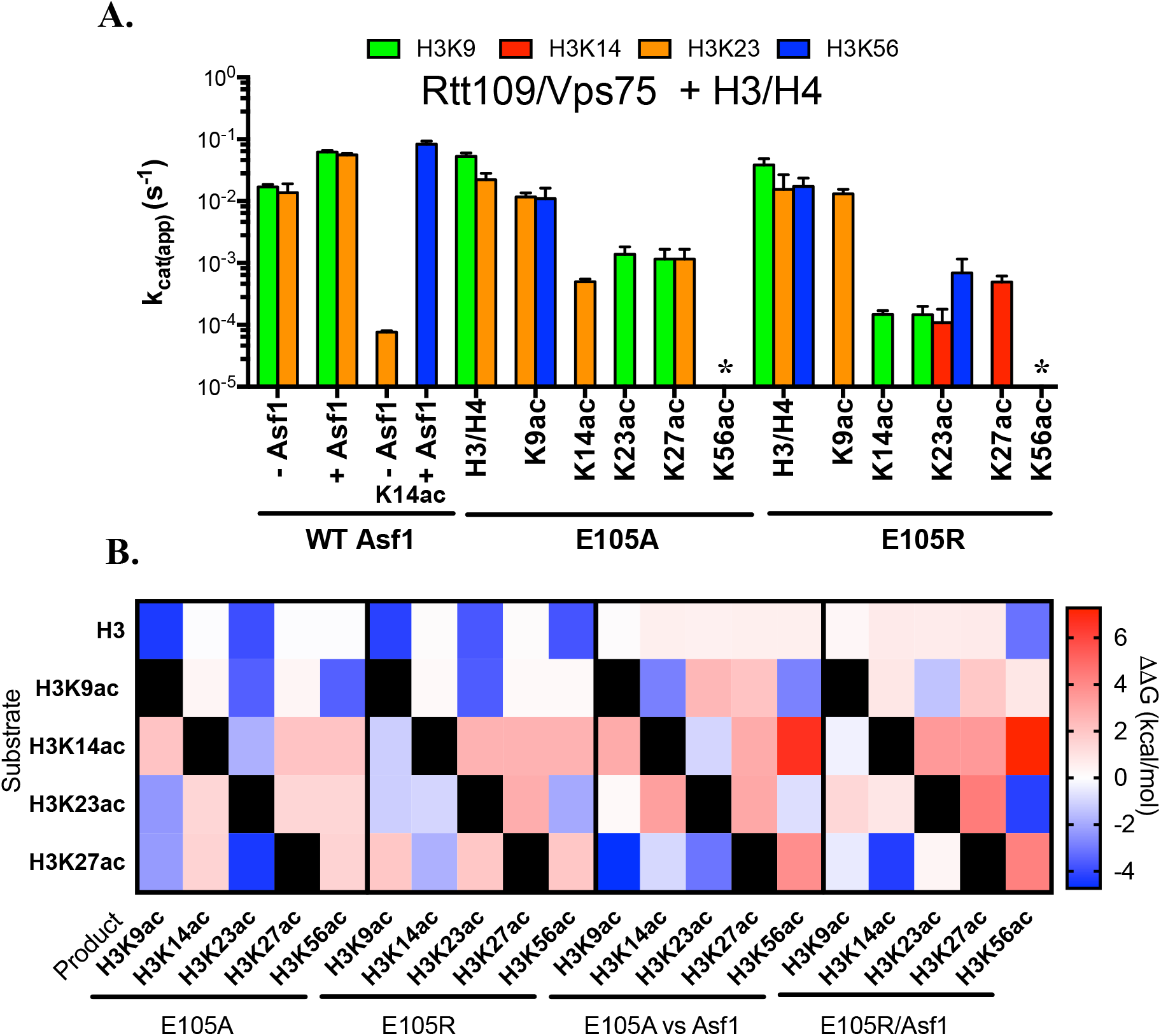
Analysis of E105A/R dependent changes to residue selectivity as a function of pre-acetylated states of histone H3. A) Comparison of site-specific k_cat(app)_ of H3/H4 (with and without singly acetylated H3) by Rtt109-Vps75 acetylation in the presence of Asf1 E105A/R mutants. The error bar represents the standard error in k_cat(app)_. The apparent k_cat_ are summarized in Table 5. B) Free energy changes to selectivity due to different acetylation states of histone H3 with E105A or R mutations to WT Asf1. Right is the free energy difference between Asf1(E105A) and Asf1 and Left is the difference between Asf1(E105R) and Asf1.

## DISCUSSION

In the present study, we have demonstrated that Asf1 can recognize PTMs to drive subsequent modifications. Specifically, Asf1 works with H3K14ac/H4 synergistically to drive H3K56 acetylation by Rtt109-Vps75. *In vitro* kinetic assays without Asf1 demonstrate that just a single acetylation can contribute to significant changes of specificity and selectivity of Rtt109-Vps75. These results reveal how pre-acetylation impacts the selectivity of subsequent acetylation sites and can explain the inconsistencies in previous studies with histones extracted from *in vivo* sources that contain various amounts of PTMs [9]. Singly acetylated H3/H4 allowed us to pinpoint how pre-acetylation influence sequential acetylation sites by Rtt109-Vps75. Furthermore, we were able to examine how histone chaperone Asf1 may additionally act as a mediator to facilitate acetylation by Rtt109-Vps75. For example, pre-acetylation at H3K23 lead to preferential acetylation by Rtt109-Vps75 at H3-K9, K14, and K27, whereas Asf1 and H3K23ac lead to acetylation at H3-K9, K14, K27, and K56. However, H3K27ac primarily drove H3-K23 and K56 acetylation and the addition of Asf1 does not change the acetylation preference. These examples show how both pre-acetylation and Asf1 can work separately and together to mediate Rtt109-Vps75 specificity and selectivity. We postulate that there is a complex network of cross-talk between histone chaperones, PTMs and KATs, where Asf1 can act as a mediator of crosstalk and facilitate the regulation of chromatin.

Although all pre-acetylated H3/H4 lead to interesting sequential acetylation selectivity, H3K14ac was especially intriguing because Rtt109-Vps75 solely drove acetylation at K56 when Asf1 was present. This suggests that previous studies using *in vivo* sources of H3/H4 were observing not Asf1 driving H3K56 acetylation, but rather Asf1 with H3K14ac driving K56 acetylation. This is consistent with genetic studies where H3K14ac facilitates DNA damage repair [32–34] and H3K56ac is believed to be a checkpoint for DNA repair activation [4, 35, 36]. Thus, H3K14ac and H3K56ac likely function in the same DNA repair pathway where H3K14ac is a precursor for H3K56ac. These findings suggest that H3K14ac may be acetylated by a different KAT, such as GCN5 [37], thus stimulating or altering the orientation of Asf1 binding to H3 tail and mediating H3K56 acetylation by Rtt109-Vps75. One interesting hypothesis is that Gcn5 acetylates H3K14 on nucleosomes, triggering a remodeling event that then opens the tetramer to Asf1 to form the H3K14ac/H4 complex that than facilities H3K56 acetylation by Rtt109-Vps75.

Asf1 to date has been thought to bind H3/H4, splitting the tetramer complex (H3/H4)2 and forming a new substrate. This effectively doubles the substrate concentration and increases the rate of acetylation by Rtt109-Vps75 [9]. In this work, Asf1 has been shown to act also as a mediator that can facilitate R11109-Vps75 selectivity. Structural analysis of Asf1 revealed an acidic patch on Asf1 that likely binds the tail of H3. Pre-acetylation will remove the positive charge on a given lysine residue, subsequently altering the Asf1-H3 tail complex and exposing distinct residues for Rtt109-Vps75 acetylation. Furthermore, this acidic patch is highly conserved between yeast Asf1 (used in the study) and Asf1A and Asf1B found in humans (Figure S2), suggesting that finding in this work may be conserved in humans.

Although there are many differences between Rtt109 and *Af*Rtt109 structures discussed earlier [20], there is now structural evidence to support our hypothesis that the H3 tail wraps around the top of Asf1 and binds to the acidic patch. Even without the first 40 amino acids of H3, it is clear from the structure that the resolved portion of the H3 tail is positioned perfectly to wrap around Asf1 and bind to the acidic patch (Figure 5). Furthermore, Asf1-H3/H4 binds Rtt109 on the opposite side of the binding site of Vps75 (Figure S3). It is possible that crystallizing the Rtt109-Vps75-Asf1-H3/H4 complex with pre-acetylated H3 would allow the interaction between Asf1 and the tail of H3 to be resolved by stabilizing the Asf1-H3 tail interaction.

Mutations made to the Asf1 acidic patch (D37, E39, D54, D58, E88, and E105) allowed for further understanding of the role of Asf1 in mediating Rtt109-Vps75 acetylation. These residues are conserved between yeast and humans and many have been found to be mutated in cancers exhibiting DNA damaging defects [38, 39]. Clearly, the conserved Asf1 acidic patch is crucial in order to maintain the histone code. Perhaps acetylation state and location can both alter the binding of the H3 tail to Asf1, leading to exposure of different lysine residues for Rtt109-Vps75 acetylation. Additionally, it is possible that mutations in Asf1 lead to weakening of H3/H4 tail binding, leading to Asf1-H3/H4 sampling multiple conformations. This explains the nonspecific acetylation by Rtt109-Vps75 for many of the Asf1 mutants with WT histone. Whereas when pre-acetylation of H3K14ac is present, the conformation of the H3 tail changes but the interaction between H3 tail and Asf1 is not weakened. Therefore, selectivity observed for Asf1 mutants with H3K14ac as a substrate was more specific.

Since driving H3K56ac has been shown to be important for DNA repair, mutations that behave like WT Asf1 with H3/H4 but no longer drive H3K56ac with H3K14ac are particularly interesting. Mutants E105A, E39R and D58R all behaved like WT Asf1 with H3/H4 as substrate, but when H3K14ac was used no longer drove exclusive H3K56 acetylation. Although E39R and D58R do not solely drive H3K56 acetylation, we do observe some H3K56ac along with H3K27 acetylation. Since these mutations behave like WT Asf1 with H3/H4 and still partially drive H3K56 acetylation, it is likely that E39 and D58 are essential for stabilizing Asf1 interactions with pre-acetylated H3/H4 to drive H3K56 acetylation. However, when H3K14ac was used as substrate, E105A no longer primarily acetylated H3K56, and only H3K23 acetylation was observed – consistent with H3K14ac without Asf1. Interestingly, mutations to E105 with pre-acetylated H3K27 no longer drove H3K23 and H3K56 acetylation. Asf1 containing the E105A mutation displays a slower turnover rate but the selectivity was retained in the absence of pre-acetylation, while E105R completely changes the selectivity to H3K14 acetylation. All singly pre-acetylated histones (except H3K9) with E105A/R had lower specificity and many primarily acetylated H3K9 and/or H3K23, suggesting that E105A (or this region) is important for Asf1 to drive both Rtt109-Vps75 selectivity and specificity.

By examining the electrostatic potential of each of the mutants we can gain insight into how each of them impact the charged surface of Asf1. Interestingly, mutations E105A and E39R have similar charge surfaces (Figure 7A and Figure S4). Residue E39 is located on a beta strand adjacent to E105, thus introducing an arginine residue in that spot lead to an increase in positive charge across the two beta strands effectively neutralizing E105 and mimicking the E105A mutation. Although both mutations do not behave identically when H3K14ac was used as substrate, these findings do suggest the importance of these residues and specifically E105 for binding to pre-acetylated histone. Moreover, mutations E105R and E39R in yeast have been reported to lead to a loss in Asf1 anti-silencing function, further implicating the importance of E105 for Asf1 function [40].

In summary, the results in this study provide insight into how different histone acetylation marks and histone chaperone concertedly or individually impact Rtt109-Vps75 histone acetylation. We also propose the mechanistic details of how Asf1 mediates the crosstalk between H3K14ac to drive H3K56ac by Rtt109-Vps75. In doing so we can now map the progression of acetylation in a decision tree (Figure 9), demonstrating the importance of Asf1 mediating crosstalk. These data suggest that there may be other residues that can influence the selectivity of Rtt109-Vps75, and influence downstream regulatory events. In a larger context these data provide a possible mechanism to explain the limited complexity of the observed combinatorial histone modification patterns seen *in vivo*.

**Figure 9:**
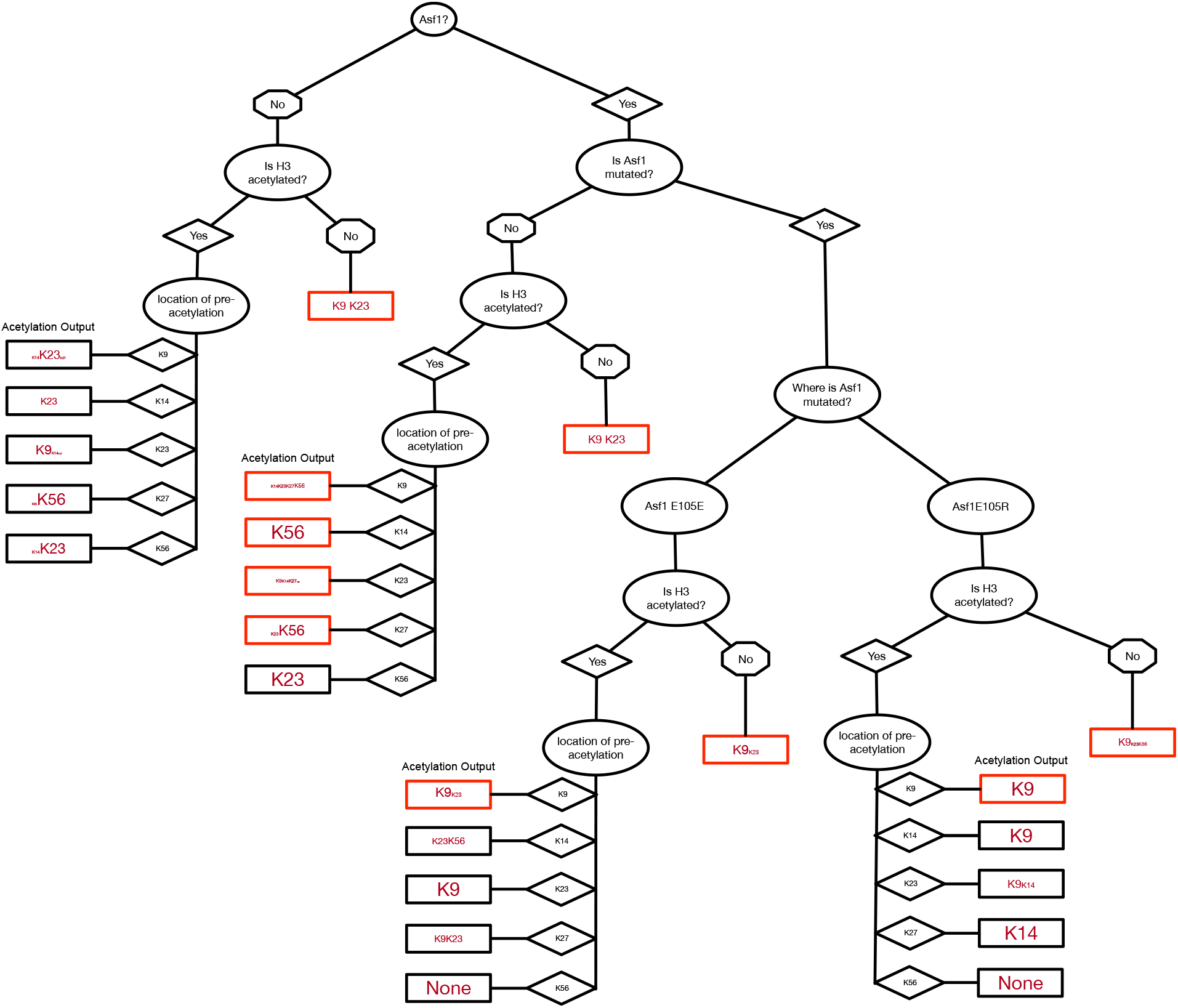
Decision tree for Asf1 dependent changes in Rtt109-Vps75 acetylation. The rectangles represent the acetylation output by Rtt109Vps75, and the difference in font represent the relative selectivity for that starting point. The red rectangles represent those substrate whose rate is as fast or faster than the acetylation of H3/H4 by Rtt109-Vps75.

## Materials and Methods

### Reagents

- All Chemicals were purchased from Sigma-Aldrich or Fisher and were either the highest commercial grade or LC/MS grade. Ultrapure water was generated from a Millipore Direct-Q 5 ultrapure water system.

### Protein preparation and purification

- All recombinant *Xenopus* histones, including WT, single lysine acetylation, uniformly ^13^C and ^15^N isotopic labeled H3, and WT H4, were purified and provided by the Protein Purification Core at Colorado State University. Histone H3/H4 tetramerization was done using previously reported methods [41]. Site-directed mutagenesis of Asf1 were purchased from GeneWiz. Rtt109-Vps75 complex, wild-type Asf1, and Asf1 mutants were expressed and purified following previous described procedures [13, 42, 43]. All purified protein concentrations were determined by UV absorbance and calculated from the extinction coefficients [44].

### k_cat_ from steady-state kinetic assays for Rtt109-Vps75

- Steady-state kinetic assays for all H3/H4 were performed under the identical buffer condition (100 mM ammonium bicarbonate and 50 mM HEPES buffer (pH 7.2)) and with saturating both H3/H4 (10 μM), Asf1 (12 μM), and acetyl-CoA (300 μM) at 37°C. The assays contained 0.04 to 3 μM Rtt109-Vps75 (varied with different substrates and histone concentrations). The initial rates (v) of acetylation were calculated from the linear increase in acetylation as a function of time prior to a total 10% acetylation. For individual lysines, the steady-state parameters k_cat_ was determined by v/[E], where [E] is the concentration of Rtt109-Vps75 and histone substrate is saturated. In our previous paper [30], we demonstrated that the ratio of the apparent k_cat_ is a valid parameter to compare the specificity for multiple substrates/sites in a multi-substrate system.

### Acetylated single-lysine H3/H4 titrations (k_cat_ conditions)

- Titrations of different acetylated H3/H4 were carried out under the same, aforementioned buffer condition in the steady-state kinetic assays. Experiments were conducted using both saturated ^13^C, ^15^N-labeled H3/H4 (10 μM) and acetyl-CoA (300 μM) at 37°C. The concentrations of singly acetylated H3/H4 were varied between 1 and 10 μM, and Rtt109-Vps75 concentrations were utilized from 0.3 to 0.45 μM, accordingly.

### Excess enzyme assays

- We performed assay under single turnover conditions to investigate the most likely acetylation spectrum on H3 of Rtt109-Vps75-mediated acetylation. The experimental condition was 1 μM H3/H4 (or any pre-acetylated H3/H4), 300 μM acetyl-CoA and 10 μM Rtt109-Vps75 in 100 mM ammonium bicarbonate and 50 mM HEPES buffer (pH 6.0) at 37°C. Buffer pH was adjusted from 7.2 to 6.0 for slowing down the acetylation reactions when Rtt109-Vps75 was excess. For all the Rtt109-Vps75 kinetics experiments, the following sample preparations, including reaction quench in ice-cold trichloroacetic acid, post-reaction propionylation using propionic anhydride, and tryptic digestion, were performed as in our previous published procedures [28].

### UPLC-MS/MS analysis

- A Waters Acquity H-class UPLC coupled with a Thermo TSQ Quantum Access triple quadrupole mass spectrometer was used to quantify the acetylated lysines on H3 peptides. The UPLC and MS/MS settings, solvent gradient, and detailed mass transitions were reported in our previously published work [37]. Retention time and specific mass transitions were both used to identify individual acetylated and propionylated peaks. The resolved peak integration was done using Xcalibur software (version 2.1, Thermo). Relative quantitative analysis was used to determine the amount of acetylation on individual lysines.

### Histone H3 acetylation with acid urea (AU) gel electrophoresis

- 30 μM H3/H4 (WT) or H3K14ac/H4, 300 μM acetyl-CoA, 0 or 36 μM Asf1 and 0 or 3 μM Rtt109-Vps75 were incubated at 37°C for 1 minute. For each experiment, at least ~0.2 μg of histones was loaded in each well of short AU gels. Short AU gels were prepared and run as previously described [45]. Histone H3 in the gels was stained by Coomassie Brilliant Blue G-250.

### Calculating the free energy of specificity and selectivity

- As we have previously shown that in a system where one substrate results in multiple products the ratio of k_cat_(s) between products is equal to the ratio of k_cat_/K_m_(s) [9, 30, 37]. This allows us to easily compare changes to specificity and selectivity by converting the ratio of k_cat_(s) to a free energy

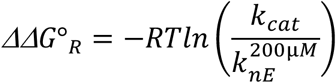

(ΔΔG°_R_) difference. We can also compare the free energy differences between k_cat_ and the non-enzymatic rate (k_nE_) at the concentration of Acetyl-CoA that is saturating for Rtt109 (200 μM) or *Pre-acetylation in the presence of Asf1 alters Rtt019-Vps75 selectivity* k_nE_^200μM^ [9]. There are residues that can be shown to be acetylated at longer times but not observed under strict steady-state conditions, where we discard all data above the time point where the total substrate (non-acetylated histone(s)) has decreased by 10% of the initial concentration. For the residues that aren’t observed under steady-state conditions we can only set the upper limit of the rate of acetylation. We set this limit by finding the fastest rate that will not produce product (<0.005%) under saturating conditions before the faster rates have consumed more than 10% of the substrate. In most cases this rate is 2-3 orders of magnitude slower than the fastest or combination

## ACKNOWLEDGEMENT

This work was supported by National Institutes of Health R01 GM102503 and a grant from the Pennsylvania Department of Health to A.J.A. J.M.C was supported by NIH Training Grant T32 CA009035-36A1.

## Supplemental Information

**Figure S1:**
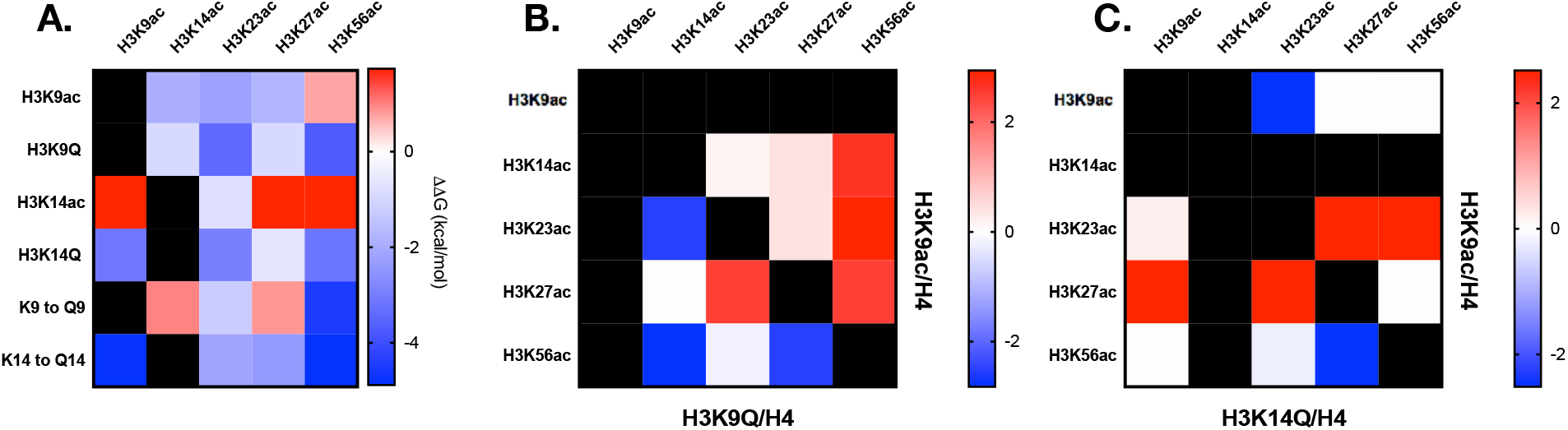
Analysis of changes to residue selectivity as a function of site specific Q mutations of histone H3 (H3K9Q and H3K14Q) with H3/H4 and pre-acetylated histone. A) Comparison of site-specific k_cat_(app) of H3/H4 with either H3K9Q or H3K914Q. H3K9ac and H3K14ac are included for comparison. K9 to Q9 and K14 to Q14 shows selectivity differences between acetylated H3 and Q mutations. B) The apparent free energy changes to selectivity due to different acetylation states of histone H3 with H3K9ac/H4 and H3K9Q/H4. C) The apparent free energy changes to selectivity due to different acetylation states of histone H3 with H3K14ac/H4 and H3K14Q/H4.

### Glutamine substitution does not act as an acetyl-lysine mimic for Rtt109-Vps75 dependent acetylation

Lysine (K) to glutamine (Q) mutations have often been used to mimic acetyl-lysines (Kac) in cells [46, 47]. Substituting a Q for K is thought to be a simple way to test the impact of a specific acetylation site in a biological context. To determine if this would be a feasible option for understanding the sequential acetylation by Rtt019-Vps75, we constructed two K to Q mutations, H3K9Q and H3K14Q. Comparing the H3K9Q to the H3K9ac it became obvious that they induced different changes to selectivity (Figure S1 and Table S1). The H3K9ac substrate was acetylated at K14, K23, and K27, whereas H3K9Q was only acetylated at K23 and K56. The H3K9Q directed the acetylation to K23 which was acetylated 5-fold faster than on the H3K9ac substrate. Unlike H3K9Q, which restricted the site(s) of acetylation, H3K14Q broadened the selectivity. Rtt109-Vps75 acetylated the H3K14Q 2-orderes of magnitude faster than H3K14ac, and in addition to acetylating K23 it also acetylated K9 and K56 at similar rates. These data suggest that K to Q mutations do not fully mimic an acetyl-lysine and that even in the absence of Asf1 they are capable of inducing H3K56ac. Consistent with these findings, there is evidence to suggest that Q is not an ideal mimic of Kac [48], and therefore, we continued to explore how histone chaperone Asf1 with pre-acetylated H3/H4 impact subsequent acetylation sites.

**Table S1.** Apparent turnover numbers (k_cat_, s^-1^) of Rtt109-Vps75 for different histone substrates

| Histone Substrate | H3K9 |  | H3K14 |  | H3K23 |  | H3K27 |  | H3K56 |  |
| --- | --- | --- | --- | --- | --- | --- | --- | --- | --- | --- |
| | $k_{\text{cat}}$ ($\text{s}^{-1}$) | $\Delta\Delta G^{\ddagger}_{\text{R}}$ (kcal/mol) | $k_{\text{cat}}$ ($\text{s}^{-1}$) | $\Delta\Delta G^{\ddagger}_{\text{R}}$ (kcal/mol) | $k_{\text{cat}}$ ($\text{s}^{-1}$) | $\Delta\Delta G^{\ddagger}_{\text{R}}$ (kcal/mol) | $k_{\text{cat}}$ ($\text{s}^{-1}$) | $\Delta\Delta G^{\ddagger}_{\text{R}}$ (kcal/mol) | $k_{\text{cat}}$ ($\text{s}^{-1}$) | $\Delta\Delta G^{\ddagger}_{\text{R}}$ (kcal/mol) |
| H3/H4 (WT)<sup>a</sup> | $(1.69 \pm 0.16) \times 10^{-2}$ | -3.74 | n.d. | n.d. | $(1.37 \pm 0.53) \times 10^{-2}$ | -3.64 | n.d. | n.d. | n.d. | n.d. |
| H3K9ac/H4 | n.a. | n.a. | $(6.16 \pm 0.93) \times 10^{-4}$ | -1.93 | $(1.83 \pm 0.19) \times 10^{-3}$ | -2.20 | $(4.59 \pm 1.03) \times 10^{-4}$ | -1.77 | n.d. | n.d. |
| H3K9Q/H4 | n.a. | n.a. | n.d. | n.d. | $(9.71 \pm 0.46) \times 10^{-3}$ | -3.44 | n.d. | n.d. | $(1.72 \pm 0.10) \times 10^{-2}$ | -3.74 |
| H3K14ac/H4 | n.d. | n.d. | n.a. | n.a. | $(7.63 \pm 0.42) \times 10^{-5}$ | -0.79 | n.d. | n.d. | n.d. | n.d. |
| H3K14Q/H4 | $(5.61 \pm 1.84) \times 10^{-3}$ | -3.14 | n.a. | n.a. | $(4.02 \pm 0.54) \times 10^{-3}$ | -2.95 | n.a. | n.a. | $(5.82 \pm 1.03) \times 10^{-3}$ | -3.16 |

<sup>a</sup>The turnover numbers for H3/H4 (WT) were previously published and are included here for comparison.

**Figure S2:**
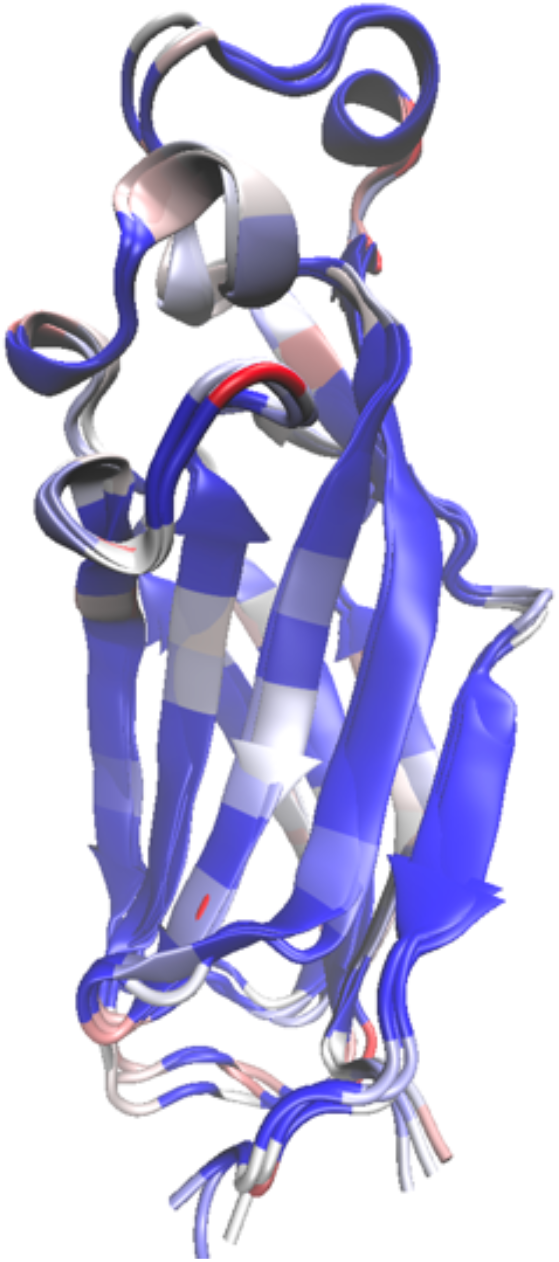
Sequence-based structural superposition was generated using Asf1 yeast (pdb:4e05) as reference structure and compared to Asf1A human (pdb: 2io5) and Asf1B human (pdb: 5bnx) structures. Alignment was done in VMD using native multiseq extension and default settings. Sequence similarity is displayed on the structure, blue represents highly conserved residues where red depicts non-conserved residues.

**Figure S3:**
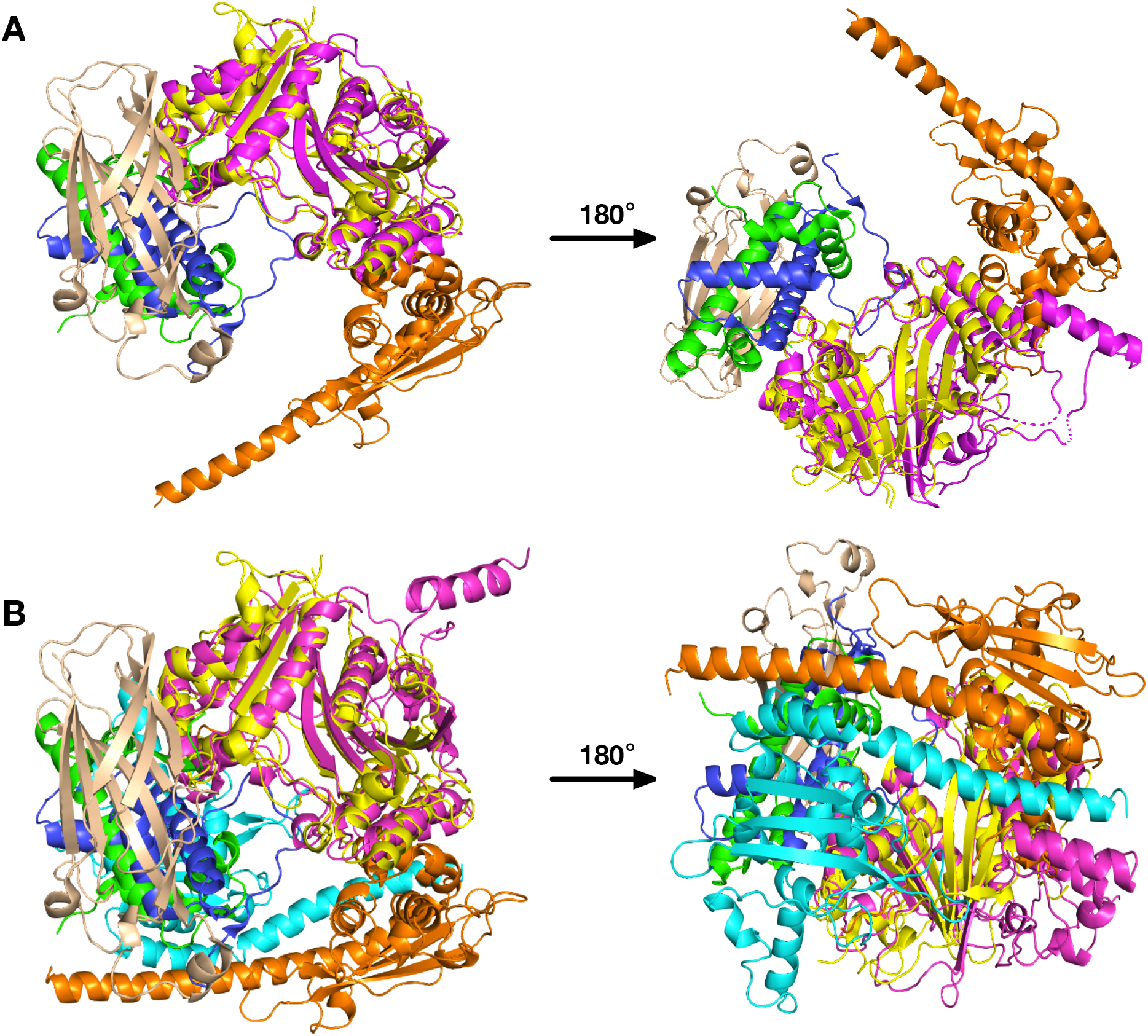
Structure alignment of *S. cerevisiae* Rtt109 (magenta) and *A. fumigatus* R1109 (yellow). A) Alignment of *A. fumigatus* R1109 in complex with Asf1 and H3/H4 (PDB: 5zba, tan and blue/green respectively) to *S. cerevisiae* Rtt109 in complex with Vps75 (orange) (PDB 3q35). B) Alignment of *A. fumigatus* R1109 in complex with Asf1 and H3/H4 (PDB: 5zba, tan and blue [H3]/green [H4] respectively) to *S. cerevisiae* Rtt109 in complex with two Vps75 (orange and cyan) (PDB 3q35).

**Figure S4:**
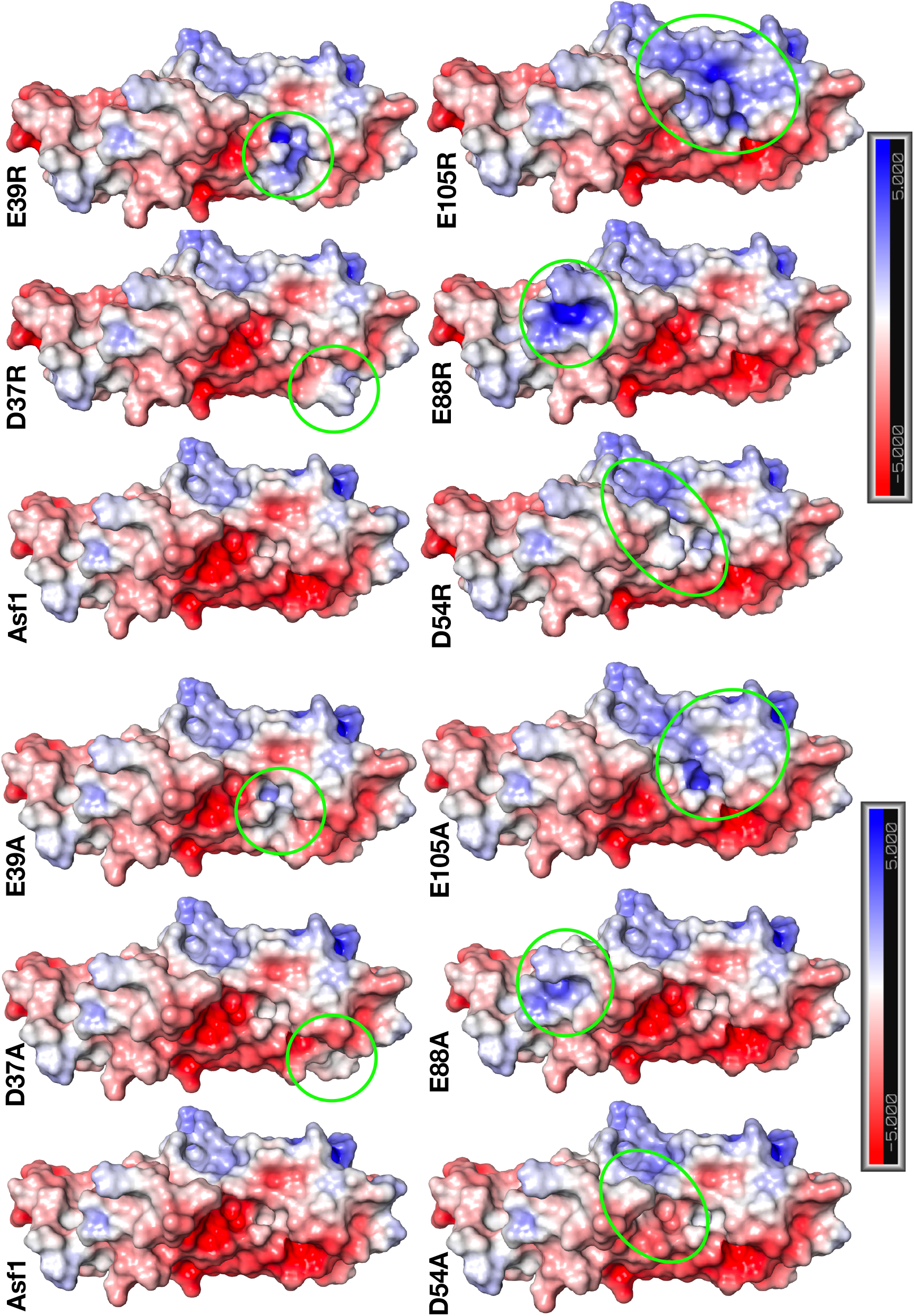
Electronic potential surface for Asf1 (PDB: 2io5), calculated using PyMOL plugin APBS electrostatics and default settings. Asf1 mutations were computationally made in PyMol using mutagenesis wizard and APBS electrostatics were calculated as for wild-type. Green circles represent the region each mutation was made.

